# An Atlas of Genetic Correlations across Human Diseases and Traits

**DOI:** 10.1101/014498

**Authors:** Brendan Bulik-Sullivan, Hilary K Finucane, Verneri Anttila, Alexander Gusev, Felix R. Day, ReproGen Consortium, Laramie Duncan, John R. B. Perry, Nick Patterson, Elise B. Robinson, Mark J. Daly, Alkes L. Price, Benjamin M. Neale

## Abstract

Identifying genetic correlations between complex traits and diseases can provide useful etiological insights and help prioritize likely causal relationships. The major challenges preventing estimation of genetic correlation from genome-wide association study (GWAS) data with current methods are the lack of availability of individual genotype data and widespread sample overlap among meta-analyses. We circumvent these difficulties by introducing a technique for estimating genetic correlation that requires only GWAS summary statistics and is not biased by sample overlap. We use our method to estimate 300 genetic correlations among 25 traits, totaling more than 1.5 million unique phenotype measurements. Our results include genetic correlations between anorexia nervosa and schizophrenia, anorexia and obesity and associations between educational attainment and several diseases. These results highlight the power of genome-wide analyses, since there currently are no genome-wide significant SNPs for anorexia nervosa and only three for educational attainment.

## Introduction

Understanding the complex relationships between human behaviours, traits and diseases is a fundamental goal of epidemiology. In the absence of randomized controlled trials and longitudinal studies, many disease risk factors are identified on the basis of population cross-sectional correlations of variables at a single time point. Such approaches can be biased by confounding and reverse causation, leading to spurious associations [1, 2]. Genetics can help elucidate cause or effect, since inherited genetic effects cannot be subject to reverse causation and are biased by a smaller list of confounders.

The first methods for testing for genetic overlap were family studies [3–7]. The disadvantage of these methods is the requirement to measure all traits on the same individuals, which scales poorly to studies of a large number of traits, especially traits that are difficult or costly to measure (e.g., low-prevalence diseases). Genome-wide association studies (GWAS) produce effect-size estimates for specific genetic variants, so it is possible to test for shared genetics by looking for correlations in effect-sizes across traits, which does not require measuring multiple traits per individual.

A widely-used technique for testing for relationships between phenotypes using GWAS data is Mendelian randomization (MR) [1, 2], which is the specialization to genetics of instrumental variables [8]. MR is effective for traits where significant associations account for a substantial fraction of heritability [9, 10]. For many complex traits, heritability is distributed over thousands of variants with small effects, and the proportion of heritability accounted for by significantly associated variants at current sample sizes is small [11]. For such traits, MR suffers from low power and weak instrument bias [8, 12].

A complementary approach is to estimate genetic correlation, a quantity that includes the effects of all SNPs, including those that do not reach genome-wide significance (Methods). Genetic correlation is also meaningful for pairs of diseases, in which case it can be interpreted as the genetic analogue of comorbidity. The two main existing techniques for estimating genetic correlation from GWAS data are restricted maximum likelihood (REML) [13–18] and polygenic scores [19, 20]. These methods have only been applied to a few traits, because they require individual genotype data, which are difficult to obtain due to informed consent limitations.

In response to these limitations, we have developed a technique for estimating genetic correlation using only GWAS summary statistics that is not biased by sample overlap. Our method, cross-trait LD Score regression, is to single trait LD Score regression [21] and is computationally very fast. We apply this method to data from 25 GWAS and report genetic correlations for 300 pairs of phenotypes, demonstrating shared genetic bases for many complex diseases and traits.

## Results

### Overview of Methods

The method presented here for estimating genetic correlation from summary statistics relies on the fact that the GWAS effect-size estimate for a given SNP incorporates the effects of all SNPs in linkage disequilibrium (LD) with that SNP [21, 22]. For a polygenic trait, SNPs with high LD will have higher *χ*^2^ statistics on average than SNPs with low LD [21]. A similar relationship holds if we replace *χ*^2^ statistics for a single study with the product of *z*-scores from two studies of traits with non-zero genetic correlation.

More precisely, under a polygenic model [13, 15], the expected value of *z*_1*j*_*z*_2*j*_ is

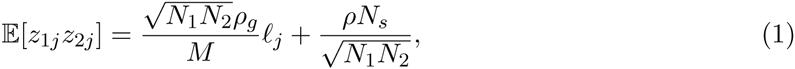

where *N_i_* is the sample size for study *i*, *ρ_g_* is genetic covariance (defined in Methods), *ℓ_j_* is LD Score [21], *N_s_* is the number of individuals included in both studies, and *ρ* is the phenotypic correlation among the *N_s_* overlapping samples. We derive this equation in the Supplementary Note. If study 1 and study 2 are the same study, then Equation 1 reduces to the single-trait result from [21], because genetic covariance between a trait and itself is heritability, and *χ*^2^ = *z*^2^. Asa consequence of equation 1, we can estimate genetic covariance using the slope from the regression of *z*_1*j*_*z*_2*j*_ on LD Score, which is computationally very fast (Methods). If there is sample overlap, it will only affect the intercept from this regression (the term 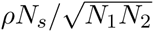) and not the slope, so the estimates of genetic correlation will not be biased by sample overlap. Similarly, shared population stratification will alter the intercept but have minimal impact on the slope, for the same reasons that population stratification has minimal impact on the slope from single-trait LD Score regression [21]. If we are willing to assume no shared population stratification and we know the amount of sample overlap and phenotypic correlation in advance (i.e., the true value of 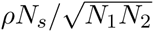), we can constrain the intercept to this value, which reduces the standard error. We refer to this approach as constrained intercept LD Score regression. Normalizing genetic covariance by the SNP-heritabilities yields genetic correlation: 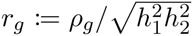, where 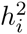 denotes the SNP-heritability [13] from study *i*. Genetic correlation ranges between −1 and 1. Similar results hold if one or both studies is a case/control study, in which case genetic covariance is on the observed scale. There is no distinction between observed and liability scale genetic correlation for case/control traits, so we can talk about genetic correlation between a case/control trait and a quantitative trait and genetic correlation between pairs of case/control traits without difficulties (Supplementary Note).

### Simulations

We performed a series of simulations to evaluate the robustness of the model to potential confounders such as sample overlap and model misspecification, and to verify the accuracy of the standard error estimates (Methods).

Table 1 shows cross-trait LD Score regression estimates and standard errors from 1,000 simulations of quantitative traits. For each simulation replicate, we generated two phenotypes for each of 2,062 individuals in our sample by drawing effect sizes approximately 600,000 SNPs on chromosome 2 from a bivariate normal distribution. We then computed summary statistics for both phenotypes and estimated heritability and genetic correlation with cross-trait LD Score regression. The summary statistics were generated from completely overlapping samples. Results are shown in Table 1. These simulations confirm that cross-trait LD Score regression yields accurate estimates of the true genetic correlation and that the standard errors match the standard deviation across simulations. Thus, cross-trait LD Score regression is not biased by sample overlap, in contrast to estimation of genetic correlation via polygenic risk scores, which is biased in the presence of sample overlap [20]. We also evaluated simulations with one quantitative trait and one case/control study and show that cross-trait LD Score regression can be applied to binary traits and is not biased by oversampling of cases (Table S1).

**Table 1:**
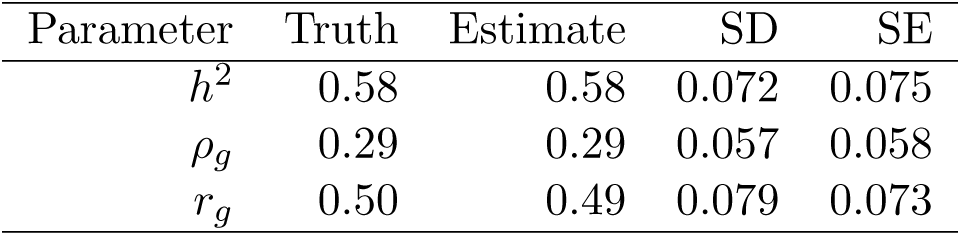
Simulations with complete sample overlap. Truth shows the true parameter values. Estimate shows the average cross-trait LD Score regression estimate across 1000 simulations. SD shows the standard deviation of the estimates across 1000 simulations, and SE shows the mean cross-trait LD Score regression SE across 1000 simulations. Further details of the simulation setup are given in the Methods.

Estimates of heritability and genetic covariance can be biased if the underlying model of genetic architecture is misspecified, *e.g.*, if variance explained is correlated with LD Score or MAF [21, 23]. Because genetic correlation is estimated as a ratio, it is more robust: biases that affect the numerator and the denominator in the same direction tend to cancel. We obtain approximately correct estimates of genetic correlation even in simulations with models of genetic architecture where our estimates of heritability and genetic covariance are biased (Table S2).

### Replication of Pyschiatric Cross-Disorder Results

As technical validation, we replicated the estimates of genetic correlations among psychiatric disorders obtained with individual genotypes and REML in [16], by applying cross-trait LD Score regression to summary statistics from the same data [24]. These summary statistics were generated from non-overlapping samples, so we applied cross-trait LD Score regression using both unconstrained and constrained intercepts (Methods). Results from these analyses are shown in Figure 1. As expected, the results from cross-trait LD Score regression were similar to the results from REML. cross-trait LD Score regression with constrained intercept gave standard errors that were only slightly larger than those from REML, while the standard errors from cross-trait LD Score regression with intercept were substantially larger, especially for traits with small sample sizes (e.g., ADHD, ASD).

**Figure 1:**
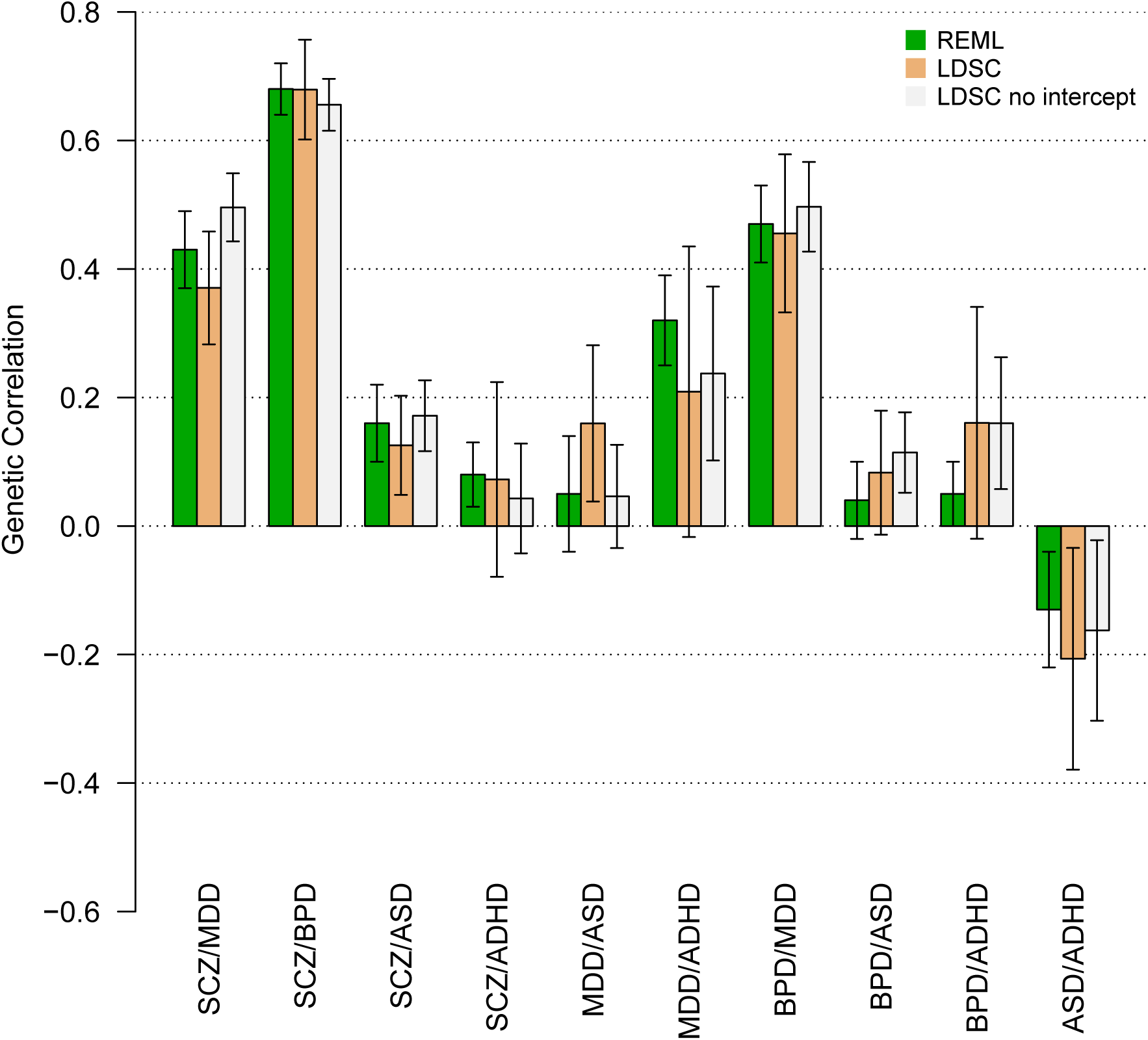
Replication of Psychiatric Cross-Disorder Results. This plot compares cross-trait LD Score regression estimates of genetic correlation using the summary statistics from [24] to estimates obtained from REML with the same data [16]. The horizontal axis indicates pairs of phenotypes, and the vertical axis indicates genetic correlation. Error bars are standard errors. Green is REML; orange is LD Score with intercept and white is LD Score with constrained intercept. The estimates of genetic correlation among psychiatric phenotypes in figure 2 use larger sample sizes; this analysis is intended as a technical validation. Abbreviations: ADHD = attention deficit disorder; ASD = autism spectrum disorder; BPD = bipolar disorder; MDD = major depressive disorder; SCZ = schizophrenia.

### Application to Summary Statistics From 25 Phenotypes

We used cross-trait LD Score regression to estimate genetic correlations among 25 phenotypes (URLs, Methods). Genetic correlation estimates for all 300 pairwise combinations of the 25 traits are shown in Figure 2. For clarity of presentation, the 25 phenotypes were restricted to contain only one phenotype from each cluster of closely related phenotypes (Methods). Genetic correlations among the educational, anthropometric, smoking, and insulin-related phenotypes that were excluded from Figure 2 are shown in Table S4 and Figures S1, S2 and S3, respectively. References and sample sizes are shown in Table S3.

**Figure 2:**
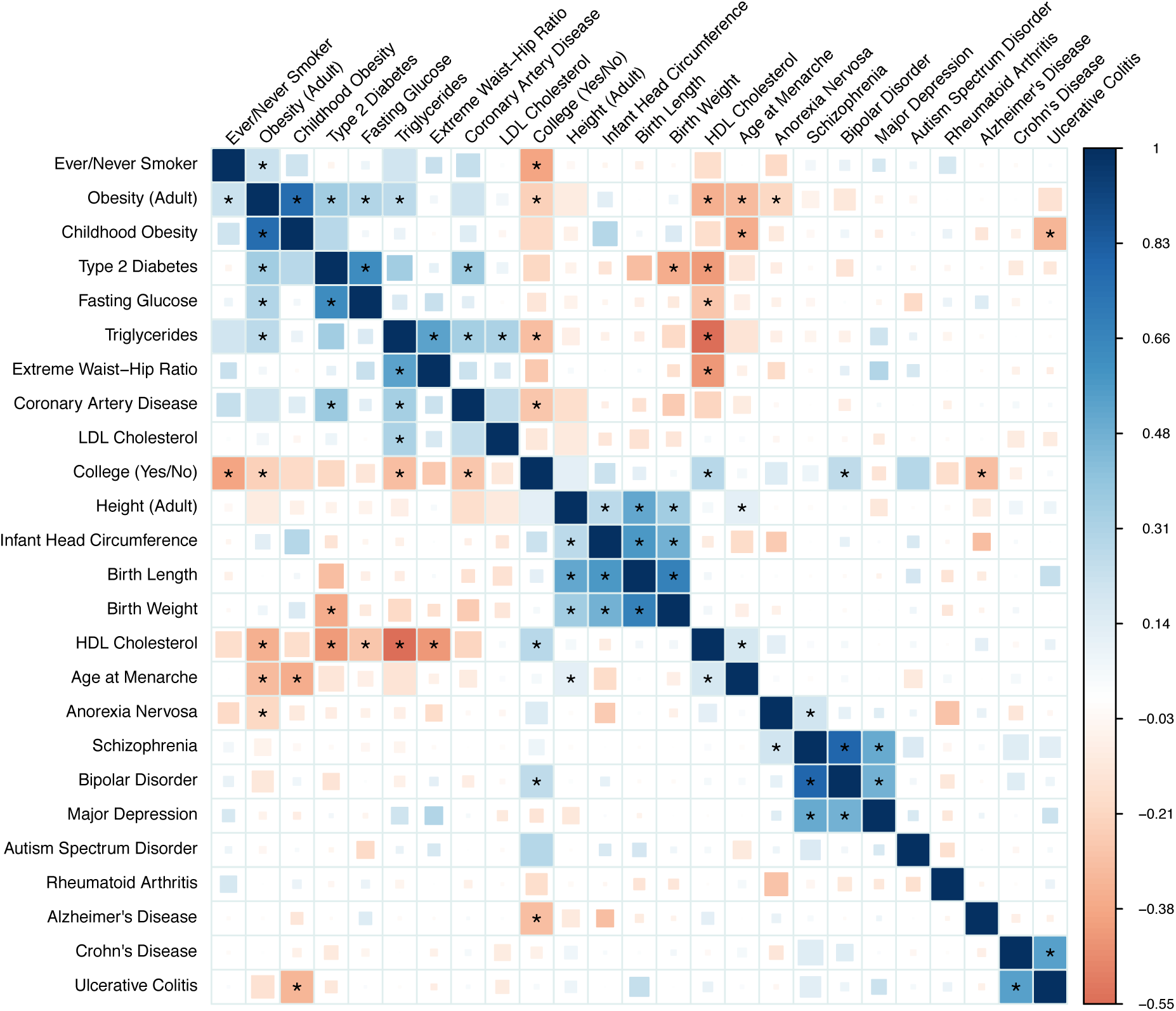
Genetic Correlations among 25 GWAS. Blue represents positive genetic correlations; red represents negative. Larger squares correspond to more significant *p*-values. Genetic correlations that are different from zero at 1% FDR are shown as full-sized squares. Genetic correlations that are significantly different from zero after Bonferroni correction for the 300 tests in this figure have an asterisk. We show results that do not pass multiple testing correction as smaller squares in order to avoid whiting out positive controls where the estimate points in the expected direction, but does not achieve statistical significance due to small sample size. This multiple testing correction is conservative, since the tests are not independent.

For the majority of pairs of traits in Figure 2, no GWAS-based genetic correlation estimate has been reported; however, many associations have been described informally based on the observation of overlap among genome-wide significant loci. Examples of genetic correlations that are consistent with overlap among top loci include the correlations between plasma lipids and cardiovascular disease [10]; age at onset of menarche and obesity [25]; type 2 diabetes, obesity, fasting glucose, plasma lipids and cardiovascular disease [26]; birth weight, adult height and type 2 diabetes [27, 28]; birth length, adult height and infant head circumference [29, 30]; and childhood obesity and adult obesity [29]. For many of these pairs of traits, we can reject the null hypothesis of zero genetic correlation with overwhelming statistical significance (*e.g.*, *p* < 10^−20^ for age at onset of menarche and obesity).

The first section of Table 2 lists genetic correlation results that are consistent with epidemiological associations, but, as far as we are aware, have not previously been reported using genetic data. The estimates of the genetic correlation between age at onset of menarche and adult height [31], triglycerides [32] and type 2 diabetes [32, 33] are consistent with the epidemiological associations.

**Table 2:** Genetic correlation estimates, standard errors and *p*-values for selected pairs of traits. Results are grouped into genetic correlations that are new genetic results, but are consistent with established epidemiological associations (“Epidemiological”), genetic correlations that are new both to genetics and epidemiology (“New/Nonzero”) and interesting null results (“New/Low”). The *p*-values are uncorrected *p*-values. Results that pass multiple testing correction for the 300 tests in Figure 2 at 1% FDR have a single asterisk; results that pass Bonferroni correction have two asterisks. We present some genetic correlations that agree with epidemiological associations but that do not pass multiple testing correction in these data.

| | Phenotype 1 | Phenotype 2 | $r_g$ (se) | $p$-value |
| --- | --- | --- | --- | --- |
| Epidemiological | Age at Menarche | Height (Adult) | 0.11 (0.03) | $6 \times 10^{-5}$ ** |
| | Age at Menarche | Type 2 Diabetes | -0.13 (0.04) | $3 \times 10^{-3}$ |
| | Age at Menarche | Triglycerides | -0.15 (0.04) | $1 \times 10^{-3}$ * |
| | Coronary Artery Disease | Age at Menarche | -0.11 (0.05) | $4 \times 10^{-2}$ |
| | Coronary Artery Disease | College (Yes/No) | -0.278 (0.07) | $1 \times 10^{-4}$ ** |
| | Coronary Artery Disease | Height (Adult) | -0.17 (0.05) | $2 \times 10^{-4}$ * |
| | Alzheimer's | College (Yes/No) | -0.30 (0.08) | $1 \times 10^{-4}$ ** |
| | Bipolar Disorder | College (Yes/No) | 0.26 (0.064) | $6 \times 10^{-5}$ ** |
| | Obesity (Adult) | College (Yes/No) | -0.23 (0.04) | $2 \times 10^{-8}$ ** |
| | Triglycerides | College (Yes/No) | -0.30 (0.04) | $5 \times 10^{-12}$ ** |
| | Anorexia Nervosa | Obesity (Adult) | -0.20 (0.04) | $4 \times 10^{-6}$ ** |
| | Ever/Never Smoker | College (Yes/No) | -0.39 (0.07) | $1 \times 10^{-9}$ ** |
| | Ever/Never Smoker | Obesity (Adult) | 0.22 (0.05) | $7 \times 10^{-5}$ ** |
| New/Nonzero | Autism Spectrum Disorder | College (Yes/No) | 0.28 (0.08) | $5 \times 10^{-4}$ * |
| | Ulcerative Colitis | Childhood Obesity | -0.33 (0.08) | $3.9 \times 10^{-5}$ ** |
| | Anorexia Nervosa | Schizophrenia | 0.19 (0.04) | $1.5 \times 10^{-5}$ ** |
| New/Low | Schizophrenia | Alzheimer's | 0.05 (0.05) | 0.58 |
|  | Schizophrenia | Ever/Never Smoker | 0.03 (0.06) | 0.26 |
|  | Schizophrenia | Triglycerides | -0.05 (0.04) | 0.21 |
|  | Schizophrenia | LDL Cholesterol | -0.02 (0.03) | 0.64 |
|  | Schizophrenia | HDL Cholesterol | 0.03 (0.04) | 0.50 |
|  | Schizophrenia | Rheumatoid Arthritis | -0.05 (0.05) | 0.38 |
|  | Crohn's Disease | Rheumatoid Arthritis | -0.02 (0.09) | 0.83 |
|  | Ulcerative Colitis | Rheumatoid Arthritis | -0.09 (0.09) | 0.33 |

The estimate of a negative genetic correlation between anorexia nervosa and obesity suggests that the same genetic factors influence normal variation in BMI as well as dysregulated BMI in psychiatric illness. This result is consistent with the observation that BMI GWAS findings implicate neuronal, rather than metabolic, cell-types and epigenetic marks [34, 35]. The negative genetic correlation between adult height and coronary artery disease agrees with a replicated epidemiological association [36–38]. We observe several significant associations with the educational attainment phenotypes from Rietveld *et al.* [39]: we estimate a statistically significant negative genetic correlation between college and Alzheimer’s disease, which agrees with epidemiological results [40, 41]. The positive genetic correlation between college and bipolar disorder is consistent with previous epidemiological reports [42, 43]. The estimate of a negative genetic correlation between smoking and college is consistent with the observed differences in smoking rates as a function of educational attainment [44].

The second section of table 2 lists three results that are, to the best of our knowledge, new both to genetics and epidemiology. One, we find a positive genetic correlation between anorexia nervosa and schizophrenia. Comorbidity between eating and psychotic disorders has not been thoroughly investigated in the psychiatric literature [45, 46], and this result raises the possibility of similarity between these classes of disease. Two, we estimate a negative genetic correlation between ulcerative colitis (UC) and childhood obesity. The relationship between premorbid BMI and ulcerative colitis is not well-understood; exploring this relationship may be a fruitful direction for further investigation. Three, we estimate a positive genetic correlation between autism spectrum disorder (ASD) and educational attainment, which itself has very high genetic correlation with IQ [39, 47, 48]. The ASD summary statistics were generated using a case-pseudocontrol study design, so this result cannot be explained by the tendency for the parents of children who receive a diagnosis of ASD to be better educated than the general population [49]. The distribution of IQ among individuals with ASD has lower mean than the general population, but with heavy tails [50] (*i.e.*, an excess of individuals with low and high IQ). There is evidence that the genetic architectures of high IQ and low IQ ASD are dissimilar [51].

The third section of table 2 lists interesting examples where the genetic correlation is close to zero with small standard error. The low genetic correlation between schizophrenia and rheumatoid arthritis is interesting because schizophrenia has been observed to be protective for rheumatoid arthritis [52], though the epidemiological effect is weak, so it is possible that there is a real genetic correlation, but it is too small for us to detect. The low genetic correlation between schizophrenia and smoking is notable because of the high prevalence of smoking among individuals with schizophrenia [53]. The low genetic correlation between schizophrenia and plasma lipid levels contrasts with a previous report of pleiotropy between schizophrenia and triglycerides [54]. Pleiotropy (unsigned) is different from genetic correlation (signed; see Methods); however, the pleiotropy reported by Andreassen, *et al.* [54] could be explained by the sensitivity of the method used to the properties of a small number of regions with strong LD, rather than trait biology (Figure S5). We estimate near-zero genetic correlation between Alzheimer’s disease and schizophrenia. The genetic correlations between Alzheimers disease and the other psychiatric traits (anorexia nervosa, bipolar, major depression, ASD) are also close to zero, but with larger standard errors, due to smaller sample sizes. This suggests that the genetic basis of Alzheimer’s disease is distinct from psychiatric conditions. Last, we estimate near zero genetic correlation between rheumatoid arthritis (RA) and both Crohn’s disease (CD) and UC. Although these diseases share many associated loci [55, 56], there appears to be no directional trend: some RA risk alleles are also risk alleles for UC and CD, but many RA risk alleles are protective for UC and CD [55], yielding near-zero genetic correlation. This example highlights the distinction between pleiotropy and genetic correlation (Methods).

Finally, the estimates of genetic correlations among metabolic traits are consistent with the estimates obtained using REML in Vattikuti *et al.* [17] (Supplementary Table S4), and are directionally consistent with the recent Mendelian randomization results from Wuertz *et al.* [57]. The estimate of 0.57 (0.074) for the genetic correlation between CD and UC is consistent with the estimate of 0.62 (0.042) from Chen *et al.* [18].

## Discussion

We have described a new method for estimating genetic correlation from GWAS summary statistics, which we applied to a dataset of GWAS summary statistics consisting of 25 traits and more than 1.5 million unique phenotype measurements. We reported several new findings that would have been difficult or impossible to obtain with existing methods, including a positive genetic correlation between anorexia nervosa and schizophrenia. Our method replicated many previously-reported GWAS-based genetic correlations, and confirmed observations of overlap among genome-wide significant SNPs, MR results and epidemiological associations.

This method is an advance for several reasons: it does not require individual genotypes, genomewide significant SNPs or LD-pruning (which loses information if causal SNPs are in LD). Our method is not biased by sample overlap and is computationally fast. Furthermore, our approach does not require measuring multiple traits on the same individuals, so it scales easily to studies of thousands of pairs of traits. These advantages allow us to estimate genetic correlation for many more pairs of phenotypes than was possible with existing methods.

The challenges in interpreting genetic correlation are similar to the challenges in MR. We highlight two difficulties. First, genetic correlation is immune to environmental confounding, but is subject to genetic confounding, analogous to confounding by pleiotropy in MR. For example, the genetic correlation between HDL and CAD in Figure 2 could result from a causal effect HDL → CAD, but could also be mediated by triglycerides (TG) [10, 58], represented graphically [59] as HDL ← G → TG → CAD, where G is the set of genetic variants with effects on both HDL and TG. Extending genetic correlation to multiple genetically correlated phenotypes is an important direction for future work [60]. Second, although genetic correlation estimates are not biased by oversampling of cases, they are affected by other forms of selection bias, such as misclassification [16].

We note several limitations of cross-trait LD Score regression as an estimator of genetic correlation. First, cross-trait LD Score regression requires larger sample sizes than methods that use individual genotypes in order to achieve equivalent standard error. Second, cross-trait LD Score regression is not currently applicable to samples from recently-admixed populations. Third, we have not investigated the potential impact of assortative mating on estimates of genetic correlation, which remains as a future direction. Fourth, methods built from polygenic models, such as cross-trait LD Score regression and REML, are most effective when applied to traits with polygenic genetic architectures. For traits where significant SNPs account for a sizable proportion of heritability, analyzing only these SNPs can be more powerful. Developing methods that make optimal use of both large-effect SNPs and diffuse polygenic signal is a direction for future research.

Despite these limitations, we believe that the cross-trait LD Score regression estimator of genetic correlation will be a useful addition to the epidemiological toolbox, since it allows for rapid screening for correlations among a diverse set of traits, without the need for measuring multiple traits on the same individuals or genome-wide significant SNPs.

## Methods

### Definition of Genetic Covariance and Correlation

All definitions refer to narrow-sense heritabilities and genetic covariances. Let *S* denote a set of *M* SNPs, let *X* denote a vector of additively (0-1-2) coded genotypes for the SNPs in *S*, and let *y*_1_ and *y*_2_ denote phenotypes. Define 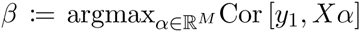, where the maximization is performed in the population (*i.e.*, in the infinite data limit). Let *γ* denote the corresponding vector for *y*_2_. This is a projection, so *β* is unique modulo SNPs in perfect LD. Define 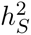, the heritability explained by SNPs in *S*, as 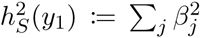 and *ρS*(*y*_1_, *y*_2_), the genetic covariance among SNPs in *S*, as 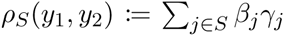. The genetic correlation among SNPs in *S* is 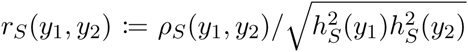, which lies in [-1,1]. Following [13], we use subscript *g* (as in 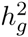, *ρ_g_*, *r_g_*) when the set of SNPs is genotyped and imputed SNPs in GWAS.

SNP genetic correlation (*r_g_*) is different from family study genetic correlation. In a family study, the relationship matrix captures information about all genetic variation, not just common SNPs. As a result, family studies estimate the total genetic correlation (*S* equals all variants). Unlike the relationship between SNP-heritability [13] and total heritability, for which 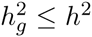, no similar relationship holds between SNP genetic correlation and total genetic correlation. If *β* and *γ* are more strongly correlated among common variants than rare variants, then the total genetic correlation will be less than the SNP genetic correlation.

Genetic correlation is (asymptotically) proportional to Mendelian randomization estimates. If we use a genetic instrument 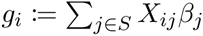 to estimate the effect *b*_12_ of *y*_1_ on *y*_2_, the 2SLS estimate is 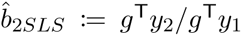 [8]. The expectations of the numerator and denominator are 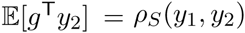 and 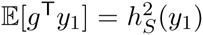. Thus, 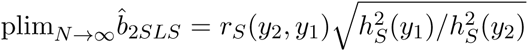. If we use the same set *S* of SNPs to estimate *b*_12_ and *b*_21_ (*e.g.*, if *S* is the set of all common SNPs, as in the genetic correlation analyses in this paper), then this procedure is symmetric in *y*_1_ and *y*_2_.

Genetic correlation is different from pleiotropy. Two traits have a pleiotropic relationship if many variants affect both. Genetic correlation is a stronger condition than pleiotropy: to exhibit genetic correlation, the directions of effect must also be consistently aligned.

### Cross-Trait LD Score Regression

We estimate genetic covariance by regressing *z*_1*j*_*z*_2*j*_ against 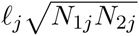, (where *N_ij_* is the sample size for SNP *j* in study *i*) then multiplying the resulting slope by *M*, the number of SNPs in the reference panel with MAF between 5% and 50% (technically, this is an estimate of *ρ*_5-50%,_ see the Supplementary Note).

If we know the amount of sample overlap ahead of time, we can reduce the standard error by constraining the intercept with the `--constrain-intercept` flag in `ldsc`. This works even if there is nonzero sample overlap, in which case the intercept should be constrained to 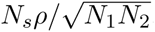.

### Regression Weights

For heritability estimation, we use the regression weights from [21]. If effect sizes for both phenotypes are drawn from a bivariate normal distribution, then the optimal regression weights for genetic covariance estimation are

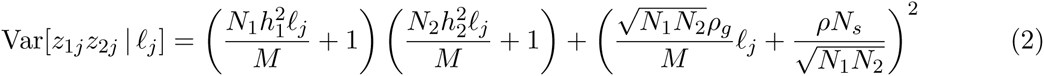

(Supplementary Note). This quantity depends on several parameters 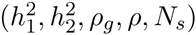 which are not known a priori, so it is necessary to estimate them from the data. We compute the weights in two steps:

1. The first regression is weighted using heritabilities from the single-trait LD Score regressions, *ρN_s_* = 0, and *ρ*_*g*_ estimated as 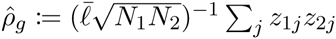.
2. The second regression is weighted using the estimates of *ρN_s_* and *ρ_g_* from step 1. The genetic covariance estimate that we report is the estimate from the second regression.

Linear regression with weights estimated from the data is called feasible generalized least squares (FGLS). FGLS has the same limiting distribution as WLS with optimal weights, so WLS *p*-values are valid for FGLS [8]. We multiply the heteroskedasticity weights by 1/*ℓ_j_* (where *ℓ_j_* is LD Score with sum over regression SNPs) in order to downweight SNPs that are overcounted. This is a heuristic: the optimal approach is to rotate the data so that it is de-correlated, but this rotation matrix is difficult to compute.

### Assessment of Statistical Significance via Block Jackknife

Summary statistics for SNPs in LD are correlated, so the OLS standard error will be biased downwards. We estimate a heteroskedasticity-and-correlation-robust standard error with a block jackknife over blocks of adjacent SNPs. This is the same procedure used in [21], and gives accurate standard errors in simulations (Table 1). We obtain a standard error for the genetic correlation by using a ratio block jackknife over SNPs. The default setting in `ldsc` is 200 blocks per genome, which can be adjusted with the `--num-blocks` flag.

### Computational Complexity

Let *N* denote sample size and *M* the number of SNPs. The computational complexity of the steps involved in LD Score regression are as follows:

1. Computing summary statistics takes 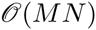 time.
2. Computing LD Scores takes 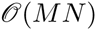 time, though the *N* for computing LD Scores need not be large. We use the *N* = 378 Europeans from 1000 Genomes.
3. LD Score regression takes 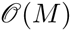 time and space.

For a user who has already computed summary statistics and downloads LD Scores from our website (URLs), the computational cost of LD Score regression is 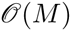 time and space. For comparison, REML takes time 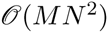 for computing the GRM and 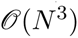 time for maximizing the likelihood.

Practically, estimating LD Scores takes roughly an hour parallelized over chromosomes, and LD Score regression takes about 15 seconds per pair of phenotypes on a 2014 MacBook Air with 1.7 GhZ Intel Core i7 processor.

### Simulations

We simulated quantitative traits under an infinitesimal model in 2062 controls from a Swedish study. To simulate the standard scenario where many causal SNPs are not genotyped, we simulated phenotypes by drawing casual SNPs from 622,146 best-guess imputed 1000 Genomes SNPs on chromosome 2, then retained only the 90,980 HM3 SNPs with MAF above 5% for LD Score regression.

We note that the simulations in [21] show that single-trait LD Score regression is only minimally biased by uncorrected population stratification and moderate ancestry mismatch between the reference panel used for estimating LD Scores and the population sampled in GWAS. In particular, LD Scores estimated from the 1000 Genomes reference panel are suitable for use with European-ancestry meta-analyses. Put another way, LD Score is only minimally correlated with *F*_*ST*_, and the differences in LD Score among European populations are not so large as to bias LD Score regression. Since we use the same LD Scores for cross-trait LD Score regression as for single-trait LD Score regression, these results extend to cross-trait LD Score regression.

### Summary Statistic Datasets

We selected traits for inclusion in the main text via the following procedure:

1. Begin with all publicly available non-sex-stratified European-only summary statistics.
2. Remove studies that do not provide signed summary statistics.
3. Remove studies not imputed to at least HapMap 2.
4. Remove studies that include heritable covariates [61].
5. Remove all traits with heritability *z*-score below 4. Genetic correlation estimates for traits with heritability *z*-score below 4 are generally too noisy to interpret.
6. Prune clusters of correlated phenotypes (*e.g.*, obesity classes 1-3) by picking the trait from each cluster with the highest heritability heritability *z*-score.

We then applied the following filters (implemented in the script `sumstats_to_chisq.py` included with `ldsc`):

1. For studies that provide a measure of imputation quality, filter to INFO above 0.9.
2. For studies that provide sample MAF, filter to sample MAF above 1%.
3. In order to restrict to well-imputed SNPs in studies that do not provide a measure of imputation quality, filter to HapMap3 [62] SNPs with 1000 Genomes EUR MAF above 5%, which tend to be well-imputed in most studies. This step should be skipped if INFO scores are available for all studies.
4. If sample size varies from SNP to SNP, remove SNPs with effective sample size less than 0.67 times the 90th percentile of sample size.
5. Remove indels and structural variants.
6. Remove strand-ambiguous SNPs.
7. Remove SNPs whose alleles do not match the alleles in 1000 Genomes.
8. Because the presence of outliers can increase the regression standard error, we also removed SNPs with extremely large effect sizes (*χ*^2^ > 80, as in [21]).

Genomic control (GC) correction at any stage biases the heritability and genetic covariance estimates downwards (see the Supplementary Note of [21]. The biases in the numerator and denominator of genetic correlation cancel exactly, so genetic correlation is not biased by GC correction. A majority of the studies analyzed in this paper used GC correction, so we do not report genetic covariance and heritability.

Data on Alzheimer’s disease were obtained from the following source:

> International Genomics of Alzheimer’s Project (IGAP) is a large two-stage study based upon genome-wide association studies (GWAS) on individuals of European ancestry. In stage 1, IGAP used genotyped and imputed data on 7,055,881 single nucleotide polymorphisms (SNPs) to meta-analyze four previously-published GWAS datasets consisting of 17,008 Alzheimer’s disease cases and 37,154 controls (The European Alzheimer’s Disease Initiative, EADI; the Alzheimer Disease Genetics Consortium, ADGC; The Cohorts for Heart and Aging Research in Genomic Epidemiology consortium, CHARGE; The Genetic and Environmental Risk in AD consortium, GERAD). In stage 2, 11,632 SNPs were genotyped and tested for association in an independent set of 8,572 Alzheimer’s disease cases and 11,312 controls. Finally, a meta-analysis was performed combining results from stages 1 and 2.

We only used stage 1 data for LD Score regression.

## URLs

1. `ldsc` software: github.com/bulik/ldsc
2. This paper: github.com/bulik/gencor_tex
3. PGC (psychiatric) summary statistics: www.med.unc.edu/pgc/downloads
4. GIANT (anthopometric) summary statistics: www.broadinstitute.org/collaboration/giant/index.php/GIANT_consortium_data_files
5. EGG (Early Growth Genetics) summary statistics: www.egg-consortium.org/
6. MAGIC (insulin, glucose) summary statistics: www.magicinvestigators.org/downloads/
7. CARDIoGRAM (coronary artery disease) summary statistics: www.cardiogramplusc4d.org
8. DIAGRAM (T2D) summary statistics: www.diagram-consortium.org
9. Rheumatoid arthritis summary statistics: www.broadinstitute.org/ftp/pub/rheumatoid_arthritis/Stahl_etal_2010NG/
10. IGAP (Alzheimers) summary statistics: www.pasteur-lille.fr/en/recherche/u744/igap/igap download. php
11. IIBDGC (inflammatory bowel disease) summary statistics: www.ibdgenetics.org/downloads.html We used a newer version of these data with 1000 Genomes imputation.
12. Plasma lipid summary statistics: www.broadinstitute.org/mpg/pubs/lipids2010/
13. SSGAC (educational attainment) summary statistics: www.ssgac.org/
14. Beans: www.barismo.com www.bluebottlecoffee.com

## Acknowledgements

We would like to thank P. Sullivan, C. Bulik, S. Caldwell, O. Andreassen for helpful comments. This work was supported by NIH grants R01 MH101244 (ALP), R03 CA173785 (HKF) and by the Fannie and John Hertz Foundation (HKF). The coffee that Brendan drank while writing this paper was roasted by Barismo in Arlington, MA and Blue Bottle Coffee in Oakland, CA.

Data on anorexia nervosa were obtained by funding from the WTCCC3 WT088827/Z/09 titled “A genome-wide association study of anorexia nervosa”.

Data on glycaemic traits have been contributed by MAGIC investigators and have been downloaded from www.magicinvestigators.org.

Data on coronary artery disease / myocardial infarction have been contributed by CARDIo-GRAMplusC4D investigators and have been downloaded from www.CARDIOGRAMPLUSC4D.ORG

We thank the International Genomics of Alzheimer’s Project (IGAP) for providing summary results data for these analyses. The investigators within IGAP contributed to the design and implementation of IGAP and/or provided data but did not participate in analysis or writing of this report. IGAP was made possible by the generous participation of the control subjects, the patients, and their families. The i-Select chips was funded by the French National Foundation on Alzheimer’s disease and related disorders. EADI was supported by the LABEX (laboratory of excellence program investment for the future) DISTALZ grant, Inserm, Institut Pasteur de Lille, Universit de Lille 2 and the Lille University Hospital. GERAD was supported by the Medical Research Council (Grant 503480), Alzheimer’s Research UK (Grant 503176), the Wellcome Trust (Grant 082604/2/07/Z) and German Federal Ministry of Education and Research (BMBF): Competence Network Dementia (CND) grant 01GI0102, 01GI0711, 01GI0420. CHARGE was partly supported by the NIH/NIA grant R01 AG033193 and the NIA AG081220 and AGES contract N01-AG-12100, the NHLBI grant R01 HL105756, the Icelandic Heart Association, and the Erasmus Medical Center and Erasmus University. ADGC was supported by the NIH/NIA grants: U01 AG032984, U24 AG021886, U01 AG016976, and the Alzheimer’s Association grant ADGC-10-196728.

## Author Contributions

MJD provided reagents. BMN and ALP provided reagents. CL, ER, VA, JP and FD aided in the interpretation of results. JP and FD provided data on age at onset of menarche. The caffeine molecule is responsible for all that is good about this manuscript. BBS and HKF are responsible for the rest. All authors revised and approved the final manuscript.

## Supplementary Note

### 1.1 Quantitative Traits

Suppose we sample two cohorts with sample sizes *N*_1_ and *N*_2_. We measure phenotype 1 in cohort 1 and phenotype 2 in cohort 2. We model phenotype vectors for each cohort as *y*_1_ = *Yβ* + *δ*, and *y*_2_ = *Zγ* + *ϵ*, where *Y* and *Z* are matrices of genotypes with columns standardized to mean zero and variance one^1^, with dimensions *N*_1_ × *M* and *N*_2_ × *M*, respectively; *β* and *γ* are vectors of per-standardized genotype effect sizes, and *δ* and *ϵ* are vectors of residuals, representing environmental effects and non-additive genetic effects. In this model, *Y* and *Z* are unobserved matrices of all SNPs, including SNPs that are not genotyped.

We treat all of *Y*, *Z*, *β*, *γ*, *δ* and *ϵ* as random. We model all of these as independent, except for *β*, *γ*, *δ*, *ϵ*. Suppose that (*β*, *γ*) has mean zero and covariance matrix^2^

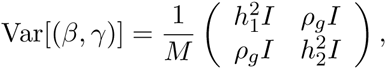

and (*δ*, *ϵ*) has mean zero and covariance matrix

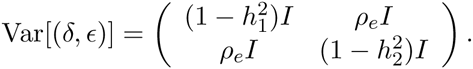

Let *ρ*:= *ρ_g_* + *ρ_e_*. Vectors of genotypes for each individual are drawn *i.i.d.* from a distribution with covariance matrix *r* (*i.e.*, *r* is an LD matrix with 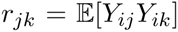). There are *N_s_* individuals who are included in both studies.

#### Lemma 1.

*Under this model, the expected genetic covariance (as defined in methods) between phenotypes is ρ_g_, justifying our use of the notation ρ_g_*.

*Proof.* Let *X* denote an 1 × *M* vector of standardized genotypes for an arbitrary individual. Under the model, the additive genetic component of phenotype1 1 for this individual is 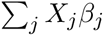, and the additive genetic component of phenotype 1 for this individual is 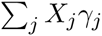. Thus, the genetic covariance between phenotype 1 and phenotype 2 is

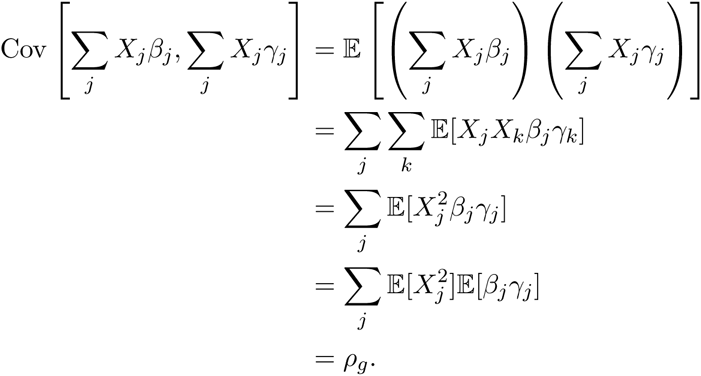

We compute linear regression *z*-scores 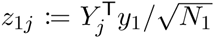 and 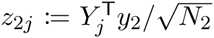 for genotyped SNPs *j* (where *Y_j_* and *Z_j_* denote the *j^th^* columns of *Y* and *Z*).

#### Definition 1.

*The LD Score of a variant j is* 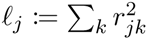, *where the sum is taken over all other variants k.*

#### Proposition 1.

*Let j denote a genotyped SNP. Under the model described above*,

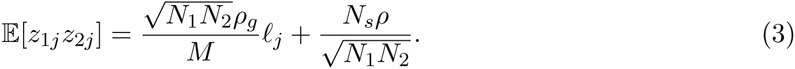

*Proof.* By the law of total expectation,

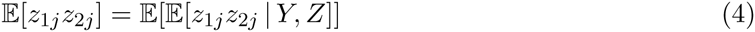

First we compute the inner expectation from Equation 4, with *Z* and *Y* fixed.

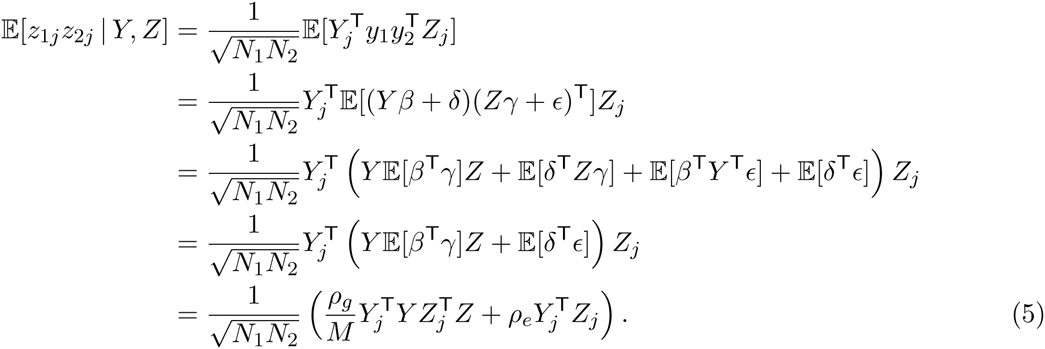

Next, we remove the conditioning on *Y* and *Z*.

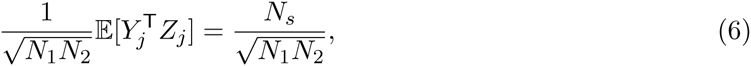

and

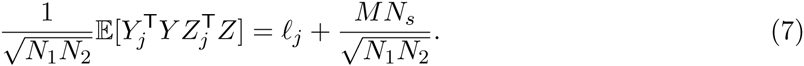

Substituting equations 6 and 7 into Equation 5,

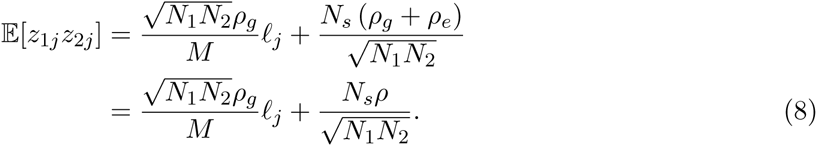

If study 1 and study 2 are the same study, then *N*_1_ = *N*_2_ = *N_s_*, 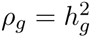 and *ρ* = 1, so Equation 8 reduces to the LD Score regression equation for a single trait from [21].

### 1.2 Regression Weights

We can improve the efficiency of LD Score regression by weighting by the reciprocal of the conditional variance function (CVF), Var[*z*_1*j*_*z*_2*j*_ | *ℓ_j_*]. The CVF is not uniquely determined by the assumptions about the first and second moments of *β* and *γ* used to derive Proposition 1. Therefore we derive the CVF for the case where *z*_1*j*_ and *z*_2*j*_ are jointly distributed as bivariate normal^3^. From a standard formula for double second moments of the bivariate normal, the CVF is

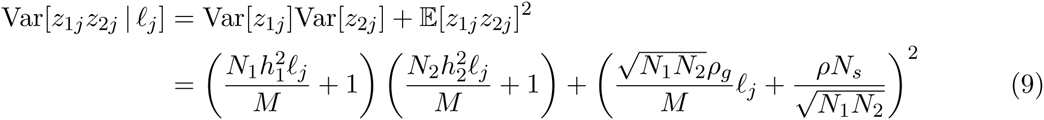

The terms on the left follow from the fact that 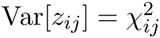 and 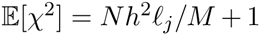. The term on the right follows from Proposition 1. Note that if *z*_1_ = *z*_2_, this reduces to the expression for the CVF of *χ*^2^ statistics from [21] (though there is an error in Equation 3.2 of the supplementary note of [21]: the right side is too small by a factor of 2. We thank Peter Visscher for pointing this out).

In cases where the normality assumption does not hold, LD Score regression will remain unbiased, but may be inefficient, because the regression weights will be suboptimal. We also apply a heuristic weighting scheme to avoid overcounting SNPs in high-LD regions, described in the methods.

### 1.3 Liability Threshold Model

In the liability threshold (probit) model [63], binary traits are determined by an unobserved continuous liability *ψ*. The observed trait is *y*:= **1**[*ψ* > *τ*], where *τ* is the liability threshold. If *ψ* is normally distributed, then setting *τ*:= Φ^−1^ (1 − *K*) (where Φ is the standard normal cdf) yields a population prevalence of *K*.

For phenotypes generated according to the liability threshold model, we can estimate not only the heritability and genetic covariance of the observed phenotype, but also the heritability and genetic covariance of the unobserved liability.

In the next lemma, we derive population case and control allele frequencies in terms of the heritability of liability when liability is generated following the model for quantitative traits from section 1.1. Since we are only modeling additive effects and are willing to assume Hardy-Weinberg equilibrium, we lose no generality and simplify notation considerably by stating the proofs in terms of haploid genotypes.

We state this lemma in terms of marginal per-allele effect sizes, instead of the per-standardized-genotype effect sizes considered in section 1.1. Here marginal means that these are the effect sizes obtained by univariate regression of phenotype against genotype in the infinite data limit. Haploid standardized genotypes are defined 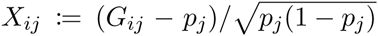, where *G_ij_* is the 0-1 coded genotype. If *β_j_* is the marginal per-standardized-genotype effect and *ζ_j_* is the marginal per-allele effect, we have *X_j_β_j_* = *G_j_ζ_j_*. Thus, setting *G_ij_* = 1 yields 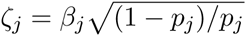.

#### Lemma 2.

*Suppose unobserved liabilities ψ*, *φ for traits y*_1_, *y*_2_ *with thresholds τ*_1_, *τ*_2_ *corresponding to prevalences K*_1_, *K*_2_ *are generated according to the mode for quantitative traits from section 1.1, i.e.*, 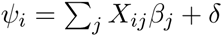, 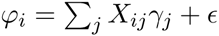, *with*

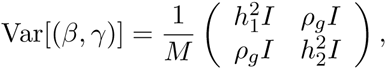

*and*

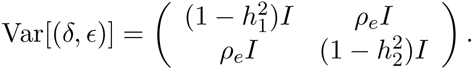

*Let ζ_j_ and ξ_j_ denote the marginal per-allele effect sizes of SNP j on ψ and φ. Let*

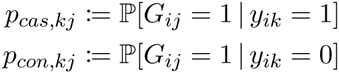

*denote the allele frequencies of SNP j in cases and controls for phenotype k, where y_ik_ denotes the value of phenotype k for individual i and k* = 1, 2*. Then*

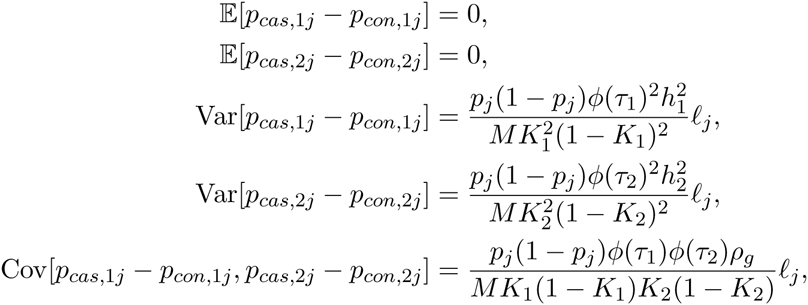

*where the expectation is taken over where φ is the standard normal density. These results apply to population allele frequencies, not allele frequencies in a finite sample. We deal with ascertained finite samples in the next section.*

*Proof.* This proof is accomplished in two steps. First, we compute allele frequencies conditional on the marginal effects on liability. To do this, we reverse the conditional probability using Bayes’ theorem, which reduces the problem to a series of [Taylor approximations to] Gaussian integrals.

Second, we remove the conditioning on the marginal effects on liability in order to express the allele frequencies in terms of 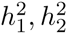, *ρ_g_* and *ℓ_j_*. Since liability is just a quantitative trait, we need only apply the LD Score regression equation for quantitative traits.

By Bayes’ rule,

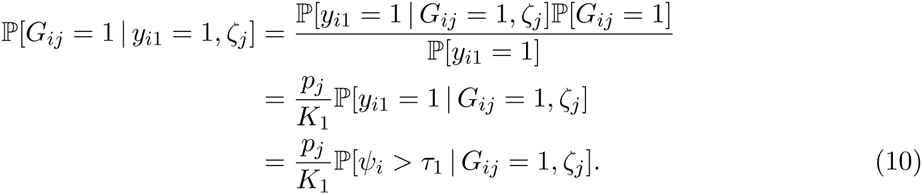

The distribution of *ψ* given *G_ij_* and *ζ_j_* is 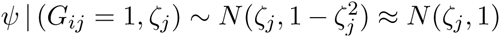 (where the approximation that the variance equals one holds when the marginal heritability explained by *j* is small, which is the typical case in GWAS). Thus 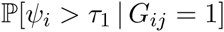 is simply a Gaussian integral. We approximate this probability with a first-order Taylor expansion around *τ*_1_.

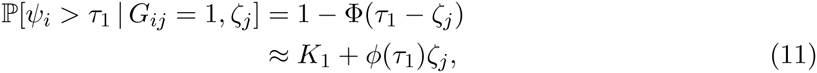

Substituting Equation 11 into Equation 10,

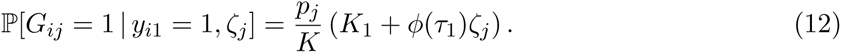

A similar argument shows that

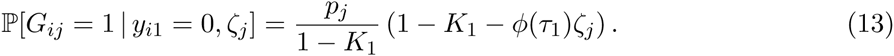

Subtracting Equation 13 from Equation 12,

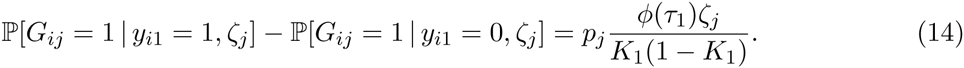

Similar results hold for trait 2, replacing *ζ* with *ξ* and subscript 1 with subscript 2.

We have written the probabilities in question in terms of constants and marginal effects on liability. Since liability is simply a quantitative trait, the means, variances, and covariances of the marginal effects on liability are described by the LD Score regression equation for quantitative traits from Proposition 1. Precisely, 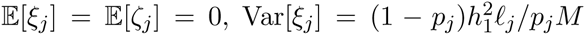, 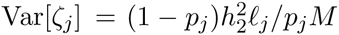 and 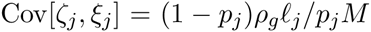. If we combine these results with Equation 14, we find that

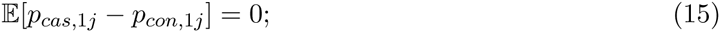

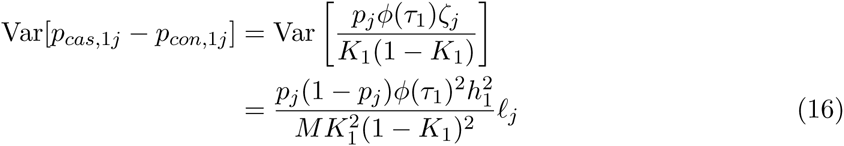

(similarly for trait two), and

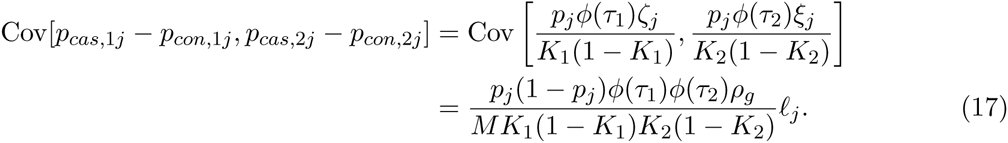

### 1.4 Ascertained Studies of Liability Threshold Traits

In the next proposition, we derive an LD Score regression equation for ascertained case/control studies.

Let *P_i_* denote the sample prevalence of *y_i_* in study *i* for *i* = 1, 2. We compute *z*-scores

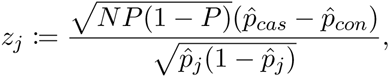

where 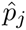 denotes allele frequency in the entire sample^4^, 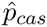 denotes sample case allele frequency and 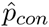 denotes sample control allele frequency.

We emphasize one subtlety before stating the main proposition. The results in this section allow for study *k* to select samples based on phenotype *l* only if *k* = *l*. If study 1 ascertains on phenotype 2 – for example, if all cases *i* in study 1 have *y_i_*_1_ = *y_i_*_2_ = 1 — then 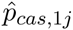 will not be an unbiased estimate of *p_cas_*_,1*j*_. Indeed, in this example, 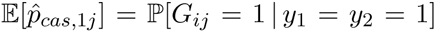, which will not equal 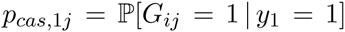 unless *ρ* = 1 or *ρ* = 0. This follows from the fact that the conditionals and marginals of a bivariate normal are equal iff *ρ* = 0 or *ρ* = 1. We do not derive formulae describing the bias, except to note that the most common scenario, the “healthy controls” model — cases are sampled independently but all controls are controls for both traits — is probably nothing to worry about, so long as cases for both traits are uncommon. In this scenario, 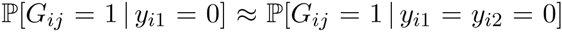. Conditioning on *y_i_*_2_ = 0 hardly changes the distribution, because *y_i_*_2_ = 0 most of the time, anyway.

#### Proposition 2.

*Under the liability threshold model from lemma 1. 3*,

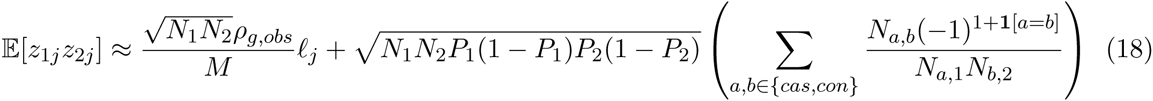

*where*

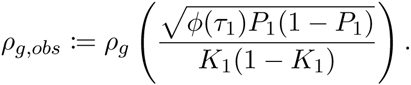

*denotes observed scale genetic covariance, N_a,b_ denotes the number of individuals with phenotype a in study 1 and b in study two for a, b* ∊ {*cas*, *con*} *(e.g., N_cas,con_ is the number of individuals who are a case in study 1 but a control in study 2), N_i_ denotes total sample size in study i and N_a,i_ for a* ∊ {*cas, con*} *and i* = 1, 2 *denotes the number of individuals with phenotype a in study i.*

Observe that 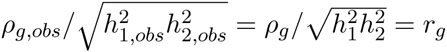. Put another way, the natural definition for “observed scale genetic correlation” turns out to be the same as regular genetic correlation, because the scale transformation factors in the numerator and denominator cancel. This is convenient: we can compute genetic correlations for binary traits on a sensible scale without having to worry about sample and population prevalences.

*Proof.* The full form of *z*_1*j*_*z*_2*j*_ is

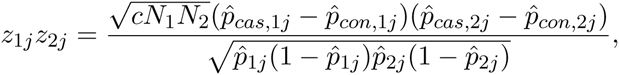

where *c*:= *P*_1_(1 − *P*_1_)*P*_2_(1 − *P*_2_). Our strategy for obtaining the expectation is

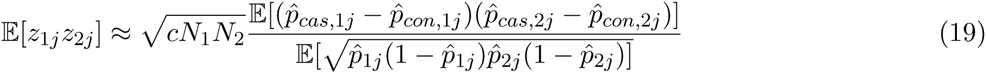

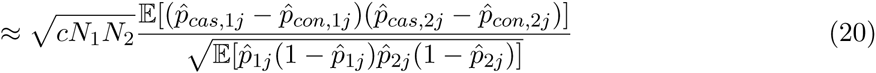

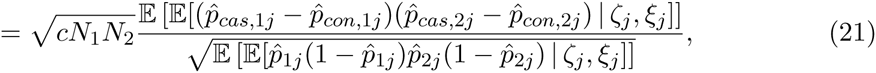

where *ζ_j_* and *ξ_j_* denote the marginal per-allele effects of *j*. Approximation 19 hides 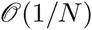 error from moving from the expectation of a ratio to a ratio of expectations. Approximation 20 hides 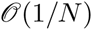 error from moving from the expectation of a square root to a square root of expectations, and dear reader we admire your perseverance in making it this far. Equality 21 follows from applying of the law of total expectation to the numerator and denominator.

First, we compute the numerator. By linearity of expectation,

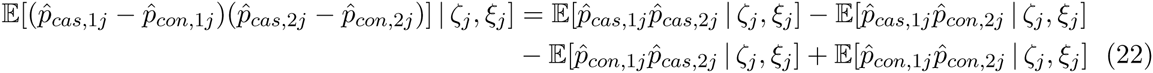

After conditioning on the marginal effects *ζ_j_* and *ξ_j_*, the only source of variance in the sample allele frequencies 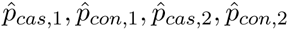 is sampling error. Write 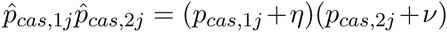, where *η* and *ν* denote sampling error. If study 1 and study 2 share samples, *ν* and *η* will be correlated:

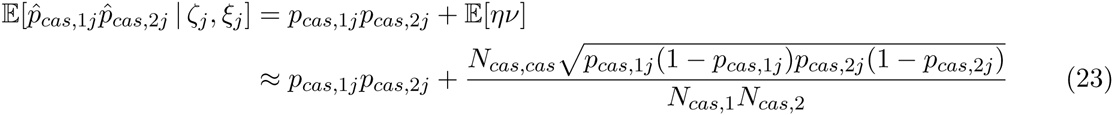

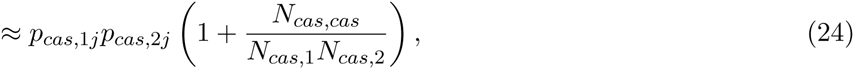

where approximation 23 is the (bivariate) central limit theorem, and approximation 24 comes from ignoring the difference between 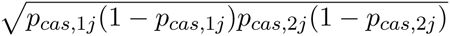 and *p_j_*(1 − *p_j_*). This step is justified in the derivation of the denominator. Similar relationships hold for the other terms in Equation 22.

If we combine equations 24 and 17, we obtain

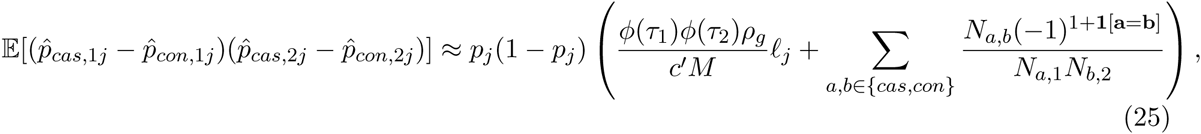

where *c*′:= *K*_1_(1 − *K*_1_)*K*_2_(1 − *K*_2_).

Next, we derive the expectation of the denominator. Conditional on *ζ_j_* and *ξ_j_*, 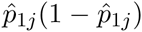 is 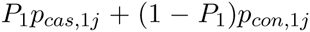 plus 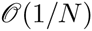 sampling variance. If studies 1 and 2 share samples, the 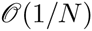 sampling variance in 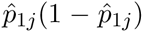 and 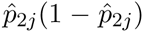 will be correlated, but this still only amounts to 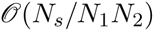 error. If we remove the conditioning on *ζ_j_* and *ξ_j_*, then 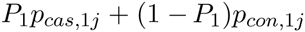 is equal to *p_j_*(1 − *p_j_*) plus 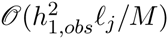 error from uncertainty in *ζ_j_*. The covariance between uncertainty in *ζ_j_* and uncertainty in *ξ_j_* is driven by *ρ_g,obs,_* so the expectation of the denominator is 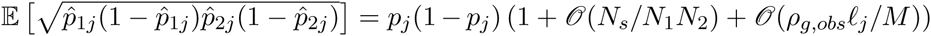. We make the approximation^5^ that

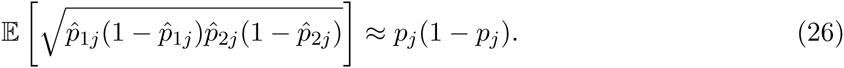

We obtain the desired result by dividing 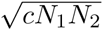 times Equation 25 by Equation 26.

#### Corollary 1.

*If study 1 is an ascertained study of a binary trait, and study 2 is a non-ascertained quantitative study, then proposition 2 holds, except with genetic covariance on the half-observed scale*

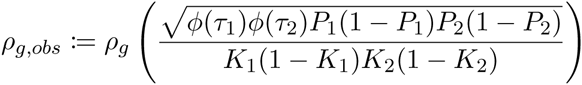

#### Corollary 2.

*For a single binary trait*,

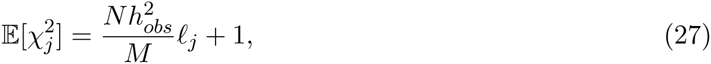

*where* 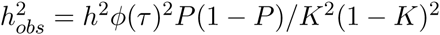.

*Proof.* This follows from proposition 2 if we set study 1 equal to study 2 and note that the observed scale genetic covariance between a trait and itself is observed scale heritability. To show that the intercept is one, observe that if study 1 and study 2 are the same, then

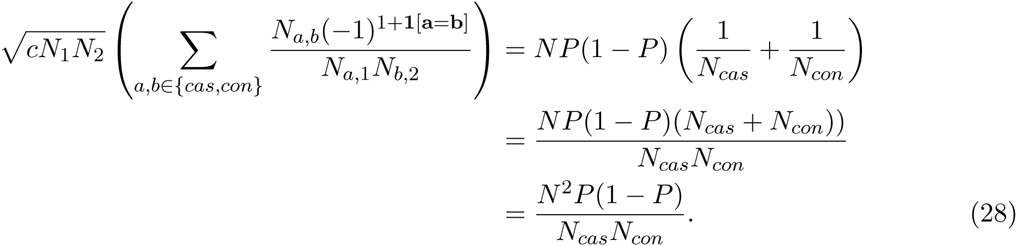

But *NP* = *N_cas_* and *N*(1 − *P*) = *N_con_*, so Equation 28 simplifies to 1.

### 1.5 Flavors of Heritability and Genetic Correlation

The heritability parameter estimated by `ldsc` is subtly different than the heritability parameter 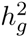 estimated by `GCTA`. If *g* denotes the set of all genotyped SNPs in some GWAS, define 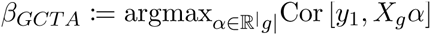, where *X_g_* is a random vector of standardized genotypes for SNPs in *g*. Then the heritability parameter estimated by `GCTA` is defined

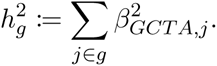

Let *S* denote the set of SNPs used to compute LD Scores (*i.e.*, 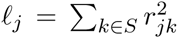), and let 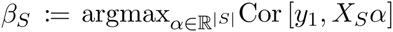. Generally *β_S,j_* ≠ *β_GCTA,j_* unless all SNPs in *S* \ *g* are not in LD with SNPs in *g*. Define

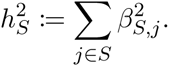

Let *S*′ denote the set of SNPs in 5 with MAF above 5%. Define

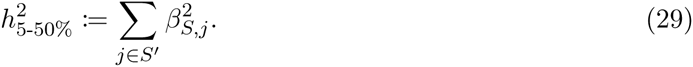

The default setting in ldsc is to report 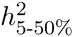, estimated as the slope from LD Score regression times *M*_5-50%,_ the number of SNPs with MAF above 5%.

The reason for this is the following: suppose that *h*^2^ per SNP is not constant as a function of MAF. Then the slope of LD Score regression will represent some sort of weighted average of the values of *h*^2^ per SNP, with more weight given to classes of SNPs that are well-represented among the regression SNPs. In a typical GWAS setting, the regression SNPs are mostly common SNPs, so multiplying the slope from LD Score regression by *M* (which includes rare SNPs) amounts to extrapolating that *h*^2^ per SNP among common variants is the same as *h*^2^ per SNP among rare variants. This extrapolation is particularly risky, because there are many more rare SNPs than common SNPs.

It is probably reasonable to treat *h*^2^ per SNP as a constant function of MAF for SNPs with MAF above 5%, but we have very little information about *h*^2^ per SNP for SNPs with MAF below 5%. Therefore we report 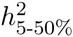 instead of 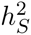 to avoid excessive extrapolation error. This lower bound can be pushed lower with larger sample sizes and better rare variant coverage, either from sequencing or imputation.

There are two main distinctions between 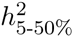 and 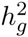. First, 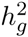 does not include the effects of common SNPs that are not tagged by the set of genotyped SNPs *g*. Second, the effects of causal 4% SNPs are not counted towards 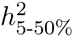. In practice, neither of these distinctions makes a large difference, since most GWAS arrays focus on common variation and manage to assay or tag almost all common variants, which is why we do not emphasize this distinction in the main text.

The relationship between the genetic covariance parameter estimated by LD Score regression and the genetic covariance parameter estimated by `GCTA` is similar to the relationship between 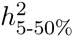 and 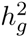. Choice of *M* is not important for genetic correlation, because the factors of *M* in the numerator and denominator cancel.

## Supplementary Tables

### Simulations with one Binary Trait and one Quantitative Trait

**Table S1:** Simulations with one binary trait and one quantitative trait. The prevalence column describes the population prevalence of the binary trait. We ran 100 simulations for each prevalence. The 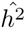 column shows the mean heritabipity estimate for the quantitative trait. The 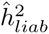 column shows the mean liability-scale heritability estimate for the binary trait. The 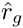 column shows the mean genetic correlation estimate. Standard deviations across 100 simulations in parentheses. The true parameter values were *r_g_* = 0.46, *h*^2^ = 0.7 for the quantitative trait and 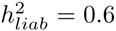 for the binary trait. For all simulations, the quantitative trait sample size was 1000, the binary trait sample size was 1000 cases and 1000 controls, and there were 500 overlapping samples. There were 1000 effective independent SNPs. The environmental covariance was 0.2. We simulated case/control ascertainment using simulated LD block genotypes and a rejection sampling model of ascertainment. This is the same strategy used to simulate case/control ascertainment in [21].

| Prevalence | $\hat{h}^2$ | $\hat{h}_{liab}^2$ | $\hat{r}_g$ |
| --- | --- | --- | --- |
| 0.01 | 0.72 (0.1) | 0.59 (0.04) | 0.51 (0.4) |
| 0.05 | 0.72 (0.12) | 0.59 (0.07) | 0.45 (0.17) |
| 0.2 | 0.72 (0.11) | 0.6 (0.08) | 0.46 (0.14) |
| 0.5 | 0.73 (0.11) | 0.59 (0.08) | 0.42 (0.17) |

### Simulations with MAF- and LD-Dependent Genetic Architecture

**Table S2:** Simulations with MAF- and LD-dependent genetic architecture. Effect sizes were drawn from normal distributions such that the variance of per-allele effect sizes was uncorrelated with MAF, and variants with LD Score below 100 were fourfold enriched for heritability. Sample size was 2062 with complete overlap between studies; causal SNPs were about 600,000 best-guess imputed 1kG SNPs on chr 2, and the SNPs retained for the LD Score regression were the subset of about 100,000 of these SNPs that were included in HM3. True parameter values are shown in the top line of the table. Estimates are averages across 100 simulations. Standard deviations (in parentheses) are standard deviations across 100 simulations. LD Scores were estimated using in-sample LD and a 1cM window. HM3 means LD Score with sum taken over SNPs in HM3. PSG (per-standardized-genotype) means LD Score with the sum taken over all SNPs in 1kG as in [21]. 30 bins means per-allele LD Score binned on a MAF by LD Score grid with MAF breaks at 0.05, 0.1, 0.2, 0.3 and 0.4 and LD Score breaks at 35, 75, 150 and 400. 60 bins means per-allele LD Score binned on a MAF by LD Score grid with MAF breaks at 0.05, 0.1, 0.15, 0.2, 0.25, 0.3, 0.35, 0.4 and 0.45 and LD Score breaks at 30, 60, 120, 200 and 300, These simulations demonstrate that naive (HM3, PSG) LD Score regression gives correct genetic correlation estimates even when heritability and genetic covariance estimates are biased, so long as genetic correlation does not depend on LD.

| LD Score | $h^2(5-50\%)$ | $\rho_g(5-50\%)$ | $r_g(5-50\%)$ |
| --- | --- | --- | --- |
| Truth | 0.83 | 0.42 | 0.5 |
| HM3 | 0.53 (0.08) | 0.28 (0.07) | 0.52 (0.1) |
| PSG | 0.36 (0.08) | 0.18 (0.06) | 0.5 (0.13) |
| 30 Bins | 0.81 (0.12) | 0.41 (0.08) | 0.51 (0.09) |
| 60 Bins | 0.81 (0.12) | 0.41 (0.09) | 0.51 (0.09) |

### Sample Sizes and References

**Table S3:** Sample sizes and references for traits analyzed in the main text.

| Trait | Reference | Sample Size |
| --- | --- | --- |
| Schizophrenia | PGC Schizophrenia Working Group, *Nature*, 2014 [64] | 70,100 |
| Bipolar disorder | PGC Bipolar Working Group, *Nat Genet*, 2011 [65] | 16,731 |
| Major depression | PGC MDD Working Group, *Mol Psych*, 2013 [66] | 18,759 |
| Anorexia Nervosa | Boraska, *et al.*, *Mol Psych*, 2014 [67] | 17,767 |
| Autism Spectrum Disorder | PGC Cross-Disorder Group, *Lancet*, 2013 [24] | 10,263 |
| Ever/Never Smoked | TAG Consortium, 2010 *Nat Genet*, [68] | 74,035 |
| Alzheimer's | Lambert, *et al.*, *Nat Genet*, 2013 [69] | 54,162 |
| College | Rietveld, *et al.*, *Science*, 2013 [39] | 101,069 |
| Height | Lango Allen, *et al.*, *Nature* 2010 [70] | 133,858 |
| Obesity Class 1 | Berndt, *et al.*, *Nat Genet*, 2013 [71] | 98,000 |
| Extreme Waist-Hip Ratio | Berndt, *et al.*, *Nat Genet*, 2013 [71] | 10,000 |
| Coronary Artery Disease | Schunkert, *et al.*, *Nat Genet*, 2011 [72] | 86,995 |
| Triglycerides | Teslovich, *et al.*, *Nature*, 2010 [73] | 96,598 |
| LDL Cholesterol | Teslovich, *et al.*, *Nature*, 2010 [73] | 95,454 |
| HDL Cholesterol | Teslovich, *et al.*, *Nature*, 2010 [73] | 99,900 |
| Type-2 Diabetes | Morris, *et al.*, *Nat Genet*, 2012 [26] | 69,033 |
| Fasting Glucose | Manning, *et al.*, *Nat Genet*, 2012 [74] | 46,186 |
| Childhood Obesity | EGG Consortium, *Nat Genet*, 2012 [29] | 13,848 |
| Birth Length | van der Valk, *et al.*, *HMG*, 2014 [75] | 22,263 |
| Birth Weight | Horikoshi, *et al.*, *Nat Genet*, 2013 [27] | 26,836 |
| Infant Head Circumference | Taal, *et al.*, *Nat Genet*, 2012 [30] | 10,767 |
| Age at Menarche | Perry, *et al.*, *Nature*, 2014 [25] | 132,989 |
| Crohn's Disease | Jostins, *et al.*, *Nature*, 2012 [76] | 20,883 |
| Ulcerative Colitis | Jostins, *et al.*, *Nature*, 2012 [76] | 27,432 |
| Rheumatoid Arthritis | Stahl, *et al.*, *Nat Genet*, 2010 [77] | 25,708 |

### Genetic Correlation between Educational Attainment Phenotypes

**Table S4:** Genetic correlation between the two educational attainment phenotypes from Rietveld, et al. [39].

| Phenotype 1 | Phenotype 2 | $r_g$ | $se$ |
| --- | --- | --- | --- |
| College (Yes/No) | Years of Education | 1.00 | 0.014 |

## Supplementary Figures

### Genetic Correlations among Anthropometric Traits

**Figure S1:**
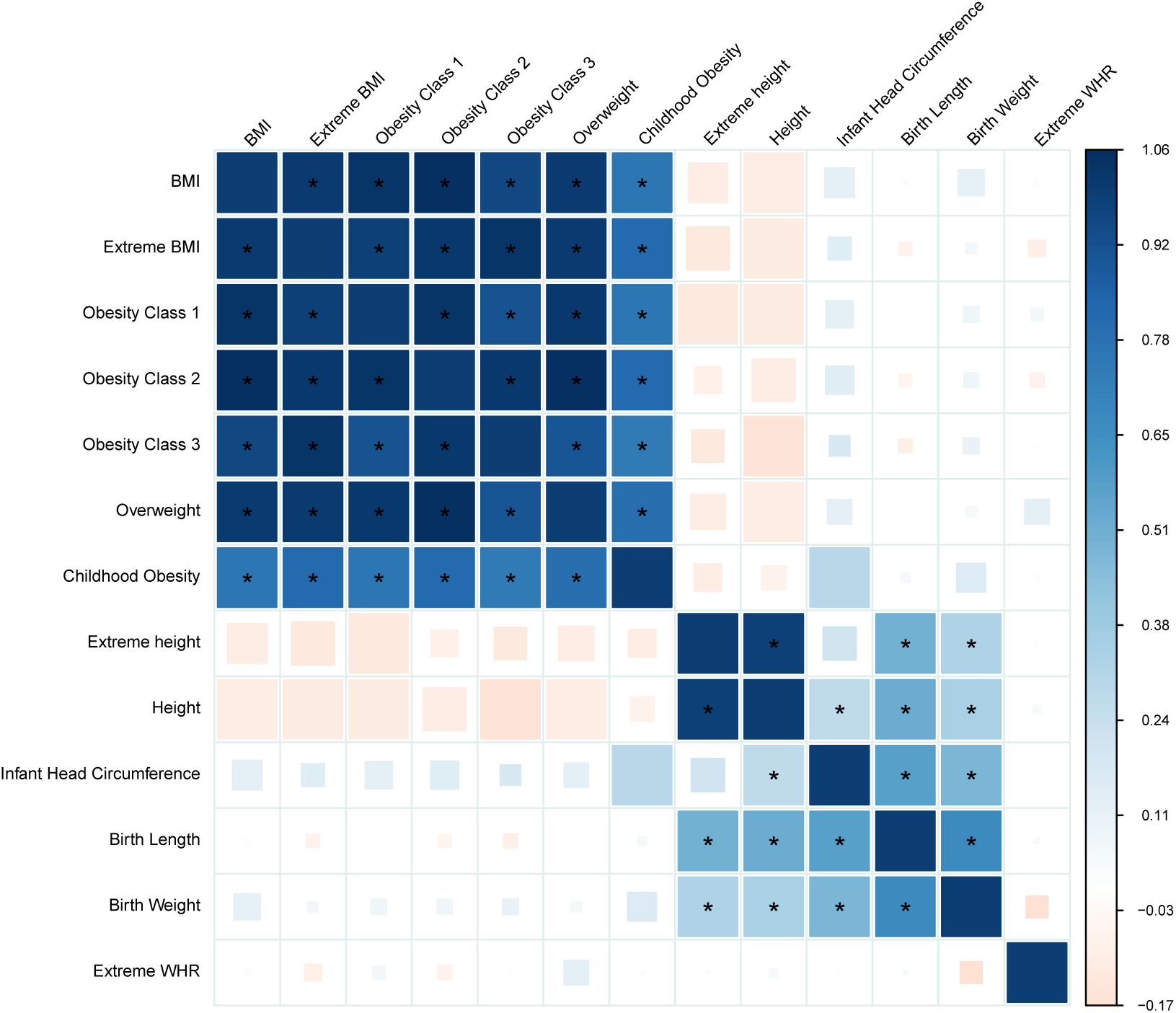
Genetic correlations among anthropometric traits from studies by the GIANT and EGG consortia. The structure of the figure is the same as Figure 2 in the main text: blue corresponds to positive genetic correlations; red corresponds to negative genetic correlation. Larger squares correspond to more significant *p*-values. Genetic correlations that are different from zero at 1% FDR are shown as full-sized squares. Genetic correlations that are significantly different from zero at significance level 0.05 after Bonferroni correction are given an asterisk.

### Genetic Correlations among Smoking Traits

**Figure S2:**
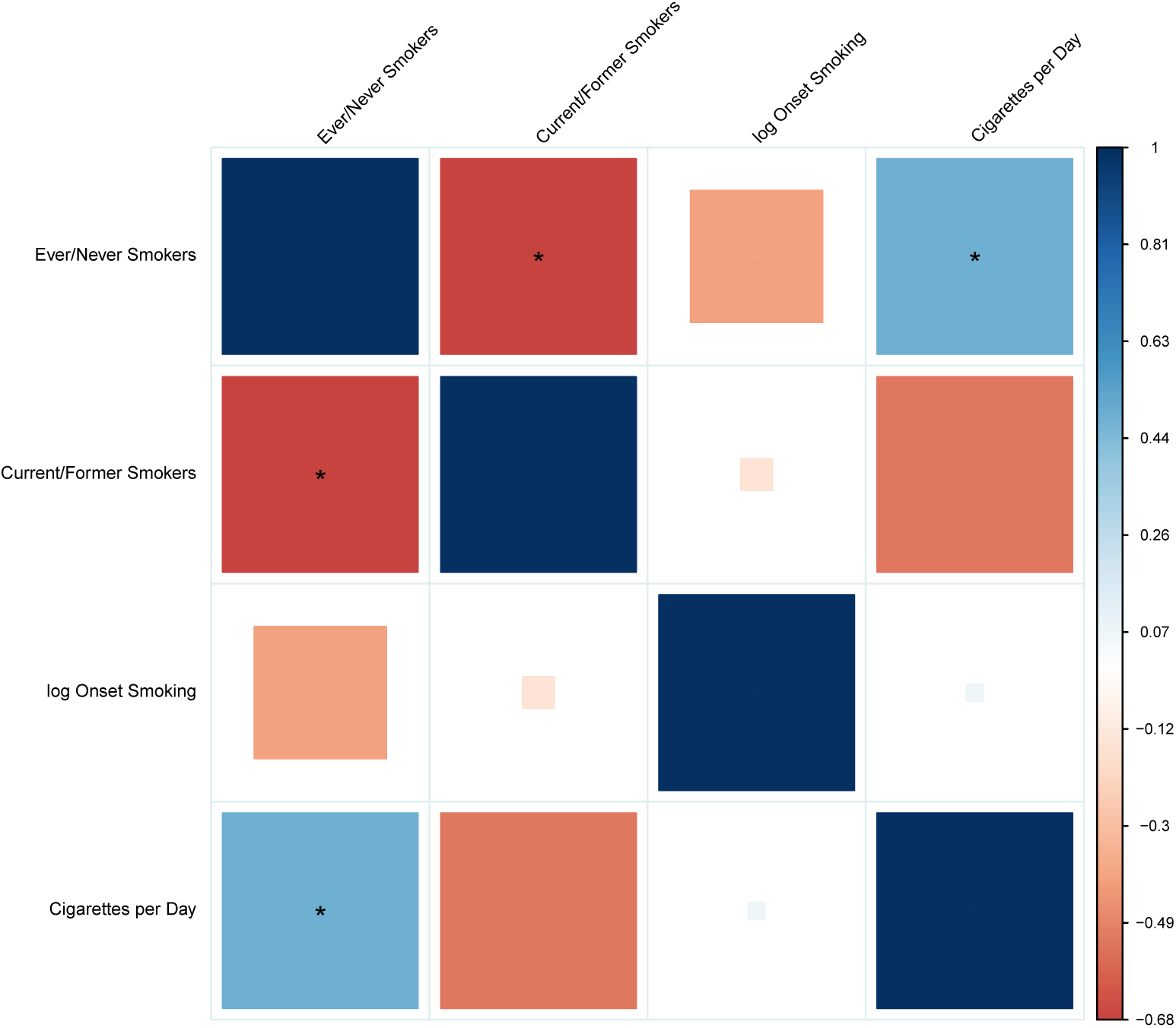
Genetic correlations among smoking traits from the Tobacco and Genetics (TAG) consortium. The structure of the figure is the same as Figure 2 in the main text: blue corresponds to positive genetic correlations; red corresponds to negative genetic correlation. Larger squares correspond to more significant *p*-values. Genetic correlations that are different from zero at 1% FDR are shown as full-sized squares. Genetic correlations that are significantly different from zero at significance level 0.05 after Bonferroni correction are given an asterisk.

### Genetic Correlations among Insulin-Related Traits

**Figure S3:**
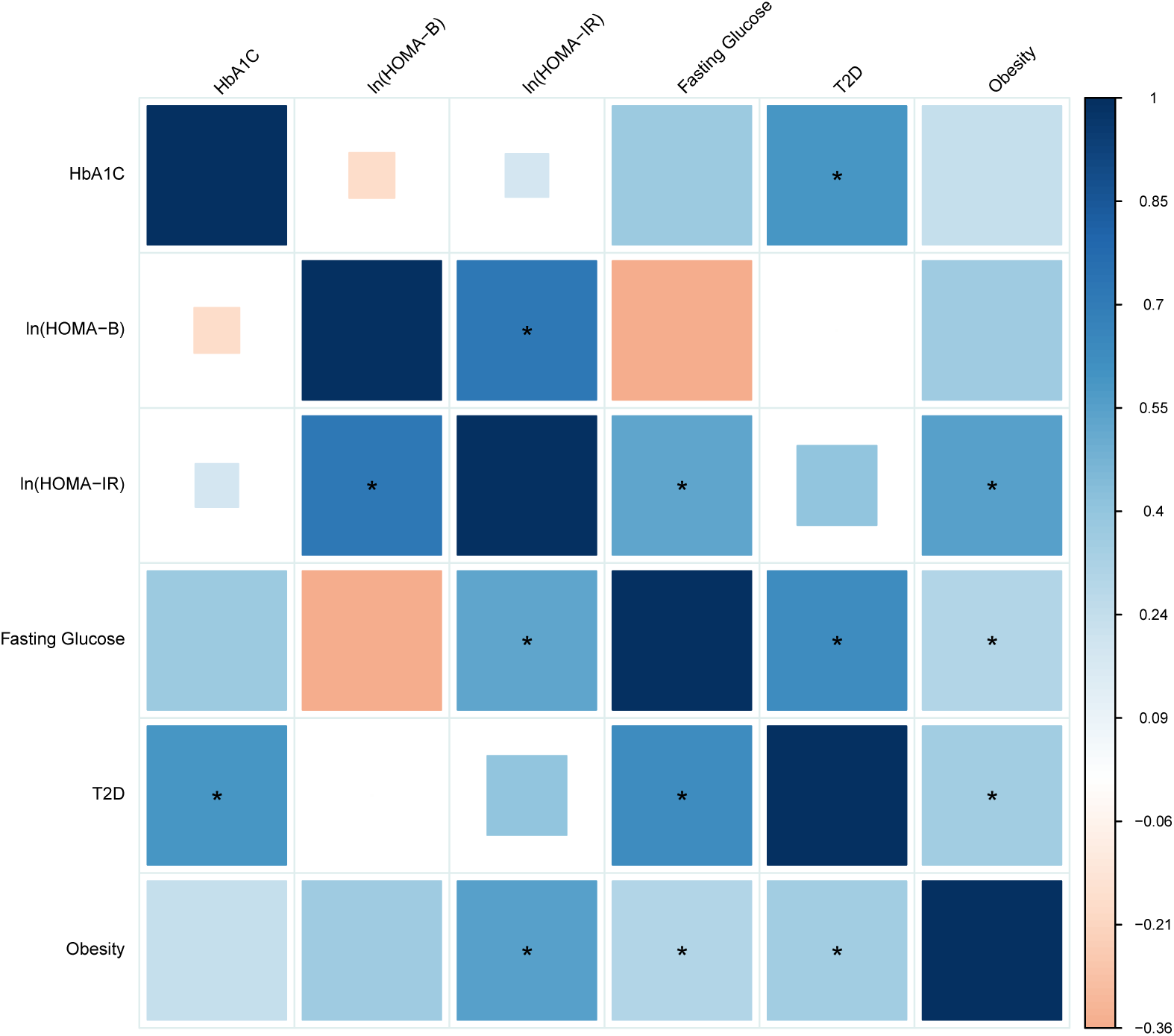
Genetic correlations among insulin-related traits from studies by the MAGIC consortium. The structure of the figure is the same as Figure 2 in the main text: blue corresponds to positive genetic correlations; red corresponds to negative genetic correlation. Larger squares correspond to more significant *p*-values. Genetic correlations that are different from zero at 1% FDR are shown as full-sized squares. Genetic correlations that are significantly different from zero at significance level 0.05 after Bonferroni correction are given an asterisk.

### Metabolic Genetic Correlations from Vattikuti, et al. and LD Score

**Figure S4:**
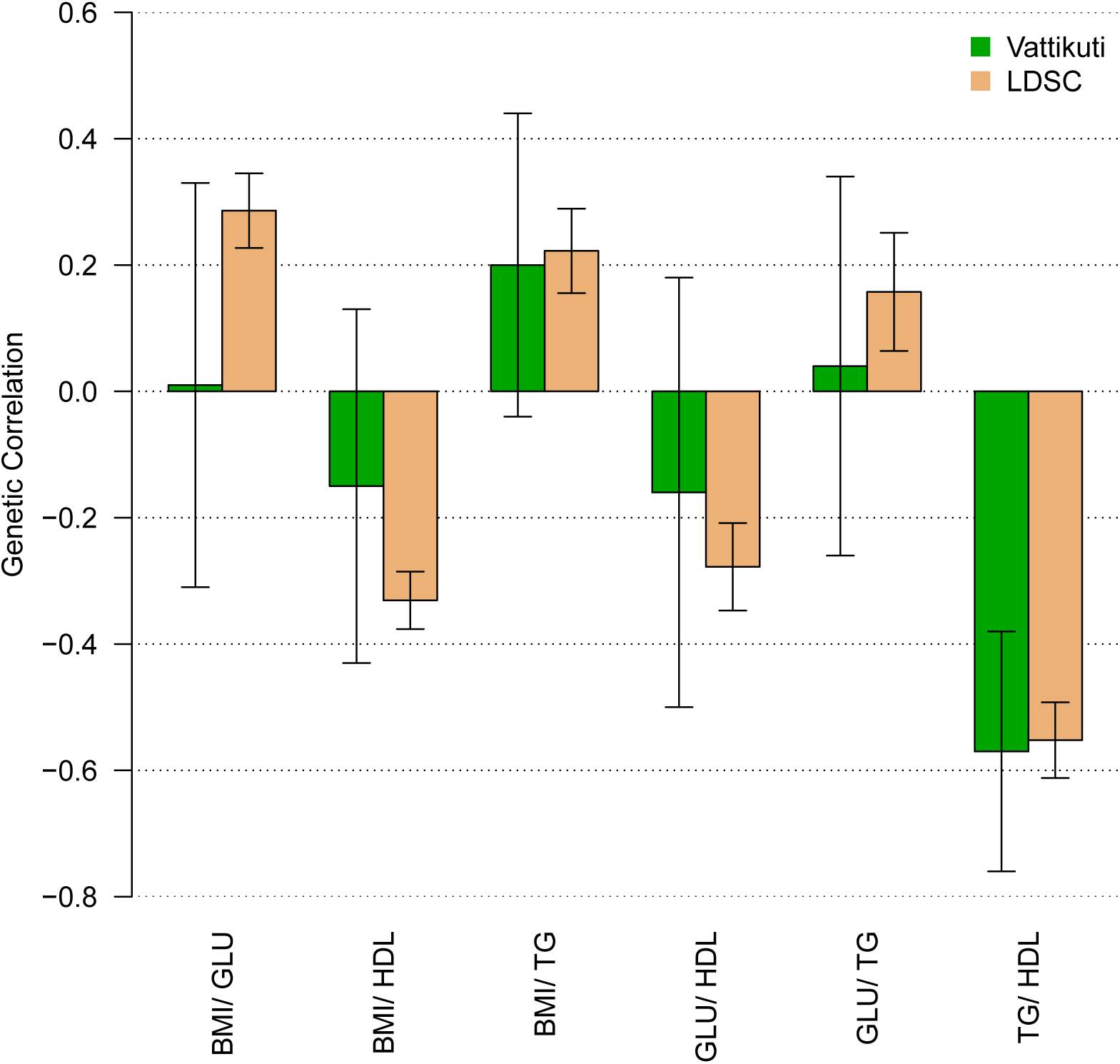
This figure compares estimates of genetic correlations among metabolic traits from table 3 of Vattikuti et al. [17] to estimates from LD Score regression. The LD Score regression estimates used much larger sample sizes. Error bars are standard errors.

### Schizophrenia — TG Conditional QQ Plot with and without the MHC

**Figure S5:**
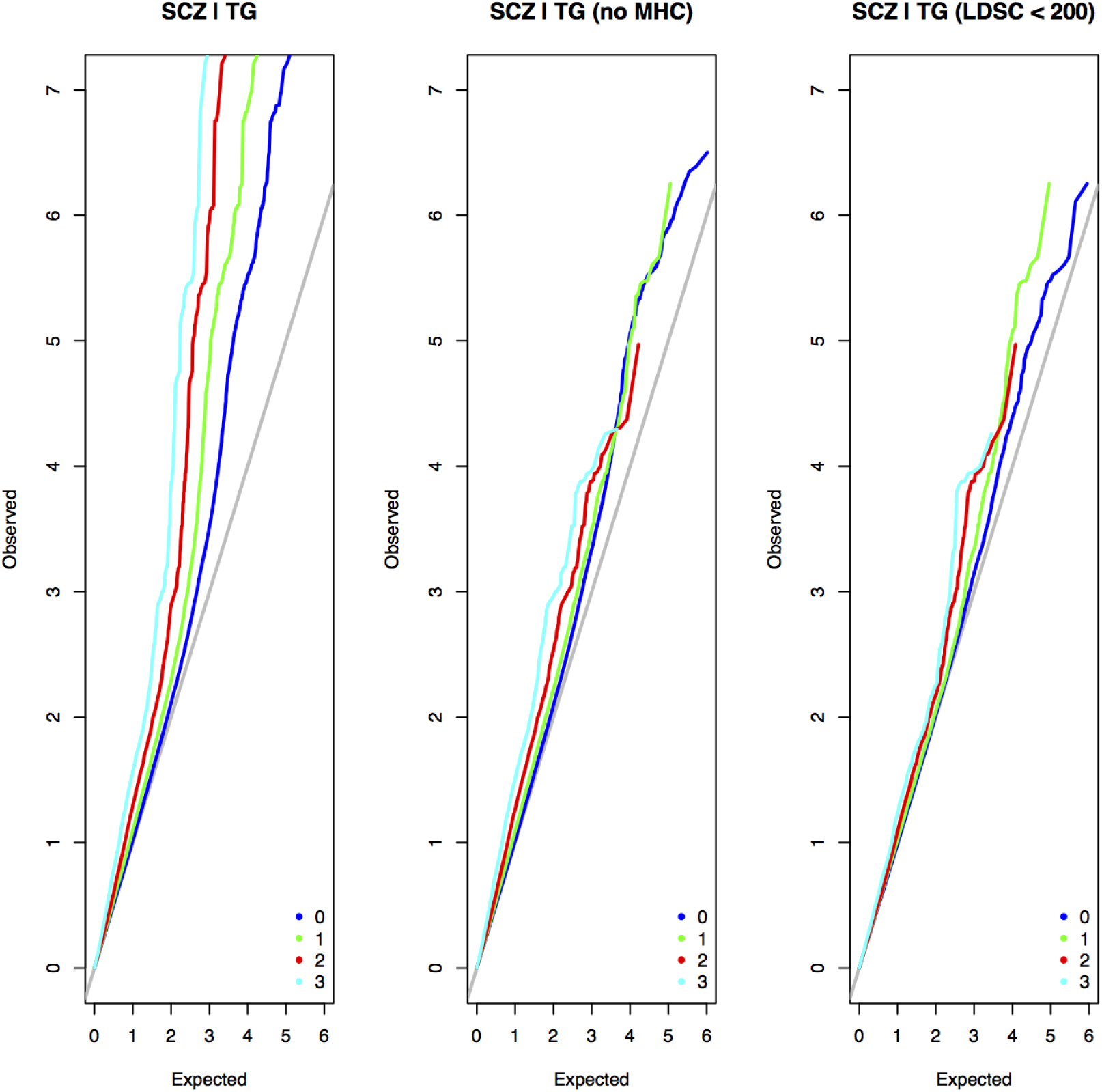
At left, we reproduced the conditional QQ plot comparing schizophrenia (SCZ) and triglycerides (TG) from Andreassen et al. [54] using the same data (PGC1 schizophrenia [78] and TG from Teslovich, et al. [73]). Conditional QQ plots show the distribution of *p*-values for SCZ conditional on the – log_10_(*p*) for TG exceeding different thresholds. The thresholds are indicated by color, as described in the legends. Dark blue corresponds to no threshold, green corresponds to – log10(*p*) > 1, red corresponds to – log10(*p*) > 2 and light blue corresponds to – log10(*p*) > 3. The major histocompatibility complex (MHC, chr6, 25-35 MB) is a genomic region containing SNPs with exceptionally long-range LD and the strongest GWAS association for schizophrenia [64], as well as an association to TG [73]. If we remove the MHC, the signal of enrichment in the conditional QQ plot is substantially attenuated (middle); in particular, the red line falls below the green and blue lines (which correspond to less stringent thresholds for TG). If in addition we remove SNPs with very high LD Scores (*ℓ* > 200, roughly the top 15% of SNPs), the signal of enrichment is further attenuated. The most likely explanation for the attenuation is that conditional QQ plots will report pleiotropy if causal SNPs are in LD (even if the causal SNPs for trait 1 are different from the casual SNPs for trait 2), which is more likely to occur in regions with long-range LD.

## Collaborators

Collaborators from the Psychiatric Genomics Consortium were, in alphabetical order: Devin Absher, Rolf Adolfsson, Ingrid Agartz, Esben Agerbo, Huda Akil, Margot Albus, Madeline Alexander, Farooq Amin, Ole A Andreassen, Adebayo Anjorin, Richard Anney, Dan Arking, Philip Asherson, Maria H Azevedo, Silviu A Bacanu, Lena Backlund, Judith A Badner, Tobias Banaschewski, Jack D Barchas, Michael R Barnes, Thomas B Barrett, Nicholas Bass, Michael Bauer, Monica Bayes, Martin Begemann, Frank Bellivier, Judit Bene, Sarah E Bergen, Thomas Bettecken, Elizabeth Bevilacqua, Joseph Biederman, Tim B Bigdeli, Elisabeth B Binder, Donald W Black, Douglas HR Blackwood, Cinnamon S Bloss, Michael Boehnke, Dorret I Boomsma, Anders D Borglum, Elvira Bramon, Gerome Breen, Rene Breuer, Richard Bruggeman, Nancy G Buccola, Randy L Buckner, Jan K Buitelaar, Brendan Bulik-Sullivan, William E Bunner, Margit Burmeister, Joseph D Buxbaum, William F Byerley, Sian Caesar, Wiepke Cahn, Guiqing Cai, Murray J Cairns, Dominique Campion, Rita M Cantor, Vaughan J Carr, Noa Carrera, Miquel Casas, Stanley V Catts, Aravinda Chakravarti, Kimberley D Chambert, Raymond CK Chan, Eric YH Chen, Ronald YL Chen, Wei Cheng, Eric FC Cheung, Siow Ann Chong, Khalid Choudhury, Sven Cichon, David St Clair, C Robert Cloninger, David Cohen, Nadine Cohen, David A Collier, Edwin Cook, Hilary Coon, Bru Cormand, Paul Cormican, Aiden Corvin, William H Coryell, Nicholas Craddock, David W Craig, Ian W Craig, Benedicto Crespo-Facorro, James J Crowley, David Curtis, Darina Czamara, Mark J Daly, Ariel Darvasi, Susmita Datta, Michael Davidson, Kenneth L Davis, Richard Day, Franziska Degenhardt, Lynn E DeLisi, Ditte Demontis, Bernie Devlin, Dimitris Dikeos, Timothy Dinan, Srdjan Djurovic, Enrico Domenici, Gary Donohoe, Alysa E Doyle, Elodie Drapeau, Jubao Duan, Frank Dudbridge, Naser Durmishi, Howard J Edenberg, Hannelore Ehrenreich, Peter Eichhammer, Amanda Elkin, Johan Eriksson, Valentina Escott-Price, Tonu Esko, Laurent Essioux, Bruno Etain, Ayman H Fanous, Stephen V Faraone, Kai-How Farh, Anne E Farmer, Martilias S Farrell, Jurgen Del Favero, Manuel A Ferreira, I Nicol Ferrier, Matthew Flickinger, Tatiana Foroud, Josef Frank, Barbara Franke, Lude Franke, Christine Fraser, Robert Freedman, Nelson B Freimer, Marion Friedl, Joseph I Friedman, Louise Frisen, Menachem Fromer, Pablo V Gejman, Giulio Genovese, Lyudmila Georgieva, Elliot S Gershon, Eco J De Geus, Ina Giegling, Michael Gill, Paola Giusti-Rodriguez, Stephanie Godard, Jacqueline I Goldstein, Vera Golimbet, Srihari Gopal, Scott D Gordon, Katherine Gordon-Smith, Jacob Gratten, Elaine K Green, Tiffany A Greenwood, Gerard Van Grootheest, Magdalena Gross, Detelina Grozeva, Weihua Guan, Hugh Gurling, Omar Gustafsson, Lieuwe de Haan, Hakon Hakonarson, Steven P Hamilton, Christian Hammer, Marian L Hamshere, Mark Hansen, Thomas F Hansen, Vahram Haroutunian, Annette M Hartmann, Martin Hautzinger, Andrew C Heath, Anjali K Henders, Frans A Henskens, Stefan Herms, Ian B Hickie, Maria Hipolito, Joel N Hirschhorn, Susanne Hoefels, Per Hoffmann, Andrea Hofman, Mads V Hollegaard, Peter A Holmans, Florian Holsboer, Witte J Hoogendijk, Jouke Jan Hottenga, David M Hougaard, Hailiang Huang, Christina M Hultman, Masashi Ikeda, Andres Ingason, Marcus Ising, Nakao Iwata, Assen V Jablensky, Stephane Jamain, Inge Joa, Edward G Jones, Ian Jones, Lisa Jones, Erik G Jonsson, Milan Macek Jr, Richard A Belliveau Jr, Antonio Julia, Tzeng Jung-Ying, Anna K Kahler, Rene S Kahn, Luba Kalaydjieva, Radhika Kandaswamy, Sena Karachanak-Yankova, Juha Karjalainen, David Kavanagh, Matthew C Keller, Brian J Kelly, John R Kelsoe, Kenneth S Kendler, James L Kennedy, Elaine Kenny, Lindsey Kent, Jimmy Lee Chee Keong, Andrey Khrunin, Yunjung Kim, George K Kirov, Janis Klovins, Jo Knight, James A Knowles, Martin A Kohli, Daniel L Koller, Bettina Konte, Ania Korszun, Robert Krasucki, Vaidutis Kucinskas, Zita Ausrele Kucinskiene, Jonna Kuntsi, Hana Kuzelova-Ptackova, Phoenix Kwan, Mikael Landen, Niklas Langstrom, Mark Lathrop, Claudine Laurent, Jacob Lawrence, William B Lawson, Marion Leboyer, Phil Hyoun Lee, S Hong Lee, Sophie E Legge, Todd Lencz, Bernard Lerer, Klaus-Peter Lesch, Douglas F Levinson, Cathryn M Lewis, Jun Li, Miaoxin Li, Qingqin S Li, Tao Li, Kung-Yee Liang, Paul Lichtenstein, Jeffrey A Lieberman, Svetlana Limborska, Danyu Lin, Chunyu Liu, Jianjun Liu, Falk W Lohoff, Jouko Lonnqvist, Sandra K Loo, Carmel M Loughland, Jan Lubinski, Susanne Lucae, Donald MacIntyre, Pamela AF Madden, Patrik KE Magnusson, Brion S Maher, Pamela B Mahon, Wolfgang Maier, Anil K Malhotra, Jacques Mallet, Sara Marsal, Nicholas G Martin, Manuel Mattheisen, Keith Matthews, Morten Mattingsdal, Robert W McCarley, Steven A McCarroll, Colm McDonald, Kevin A McGhee, James J McGough, Patrick J McGrath, Peter McGuffin, Melvin G McInnis, Andrew M McIntosh, Rebecca McKinney, Alan W McLean, Francis J McMahon, Andrew McQuillin, Helena Medeiros, Sarah E Medland, Sandra Meier, Carin J Meijer, Bela Melegh, Ingrid Melle, Fan Meng, Raquelle I Mesholam-Gately, Andres Metspalu, Patricia T Michie, Christel M Middeldorp, Lefkos Middleton, Lili Milani, Vihra Milanova, Philip B Mitchell, Younes Mokrab, Grant W Montgomery, Jennifer L Moran, Gunnar Morken, Derek W Morris, Ole Mors, Preben B Mortensen, Valentina Moskvina, Bryan J Mowry, Pierandrea Muglia, Thomas W Muehleisen, Walter J Muir, Bertram Mueller-Myhsok, Kieran C Murphy, Robin M Murray, Richard M Myers, Inez Myin-Germeys, Benjamin M Neale, Michael C Neale, Mari Nelis, Stan F Nelson, Igor Nenadic, Deborah A Nertney, Gerald Nestadt, Kristin K Nicodemus, Caroline M Nievergelt, Liene Nikitina-Zake, Ivan Nikolov, Vishwajit Nimgaonkar, Laura Nisenbaum, Willem A Nolen, Annelie Nordin, Markus M Noethen, John I Nurnberger, Evaristus A Nwulia, Dale R Nyholt, Eadbhard O’Callaghan, Michael C O’Donovan, Colm O’Dushlaine, F Anthony O’Neill, Robert D Oades, Sang-Yun Oh, Ann Olincy, Line Olsen, Edwin JCG van den Oord, Roel A Ophoff, Jim Van Os, Urban Osby, Hogni Oskarsson, Michael J Owen, Aarno Palotie, Christos Pantelis, George N Papadimitriou, Sergi Papiol, Elena Parkhomenko, Carlos N Pato, Michele T Pato, Tiina Paunio, Milica Pejovic-Milovancevic, Brenda P Penninx, Michele L Pergadia, Diana O Perkins, Roy H Perlis, Tune H Pers, Tracey L Petryshen, Hannes Petursson, Benjamin S Pickard, Olli Pietilainen, Jonathan Pimm, Joseph Piven, Andrew J Pocklington, Porgeir Porgeirsson, Danielle Posthuma, James B Potash, John Powell, Alkes Price, Peter Propping, Ann E Pulver, Shaun M Purcell, Vinay Puri, Digby Quested, Emma M Quinn, Josep Antoni Ramos-Quiroga, Henrik B Rasmussen, Soumya Raychaudhuri, Karola Rehnstrom, Abraham Reichenberg, Andreas Reif, Mark A Reimers, Marta Ribases, John Rice, Alexander L Richards, Marcella Rietschel, Brien P Riley, Stephan Ripke, Joshua L Roffman, Lizzy Rossin, Aribert Rothenberger, Guy Rouleau, Panos Roussos, Douglas M Ruderfer, Dan Rujescu, Veikko Salomaa, Alan R Sanders, Susan Santangelo, Russell Schachar, Ulrich Schall, Martin Schalling, Alan F Schatzberg, William A Scheftner, Gerard Schellenberg, Peter R Schofield, Nicholas J Schork, Christian R Schubert, Thomas G Schulze, Johannes Schumacher, Sibylle G Schwab, Markus M Schwarz, Edward M Scolnick, Laura J Scott, Rodney J Scott, Larry J Seidman, Pak C Sham, Jianxin Shi, Paul D Shilling, Stanley I Shyn, Engilbert Sigurdsson, Teimuraz Silagadze, Jeremy M Silverman, Kang Sim, Pamela Sklar, Susan L Slager, Petr Slominsky, Susan L Smalley, Johannes H Smit, Erin N Smith, Jordan W Smoller, Hon-Cheong So, Erik Soderman, Edmund Sonuga-Barke, Chris C A Spencer, Eli A Stahl, Matthew State, Hreinn Stefansson, Kari Stefansson, Michael Steffens, Stacy Steinberg, Hans-Christoph Stein-hausen, Elisabeth Stogmann, Richard E Straub, John Strauss, Eric Strengman, Jana Strohmaier, T Scott Stroup, Mythily Subramaniam, Patrick F Sullivan, James Sutcliffe, Jaana Suvisaari, Dragan M Svrakic, Jin P Szatkiewicz, Peter Szatmari, Szabocls Szelinger, Anita Thapar, Srinivasa Thirumalai, Robert C Thompson, Draga Toncheva, Paul A Tooney, Sarah Tosato, Federica Tozzi, Jens Treutlein, Manfred Uhr, Juha Veijola, Veronica Vieland, John B Vincent, Peter M Visscher, John Waddington, Dermot Walsh, James TR Walters, Dai Wang, Qiang Wang, Stanley J Watson, Bradley T Webb, Daniel R Weinberger, Mark Weiser, Myrna M Weissman, Jens R Wendland, Thomas Werge, Thomas F Wienker, Dieter B Wildenauer, Gonneke Willemsen, Nigel M Williams, Stephanie Williams, Richard Williamson, Stephanie H Witt, Aaron R Wolen, Emily HM Wong, Brandon K Wormley, Naomi R Wray, Adam Wright, Jing Qin Wu, Hualin Simon Xi, Wei Xu, Allan H Young, Clement C Zai, Stan Zammit, Peter P Zandi, Peng Zhang, Xuebin Zheng, Fritz Zimprich, Frans G Zitman, and Sebastian Zoellner.

Genetic Consortium for Anorexia Nervosa (GCAN): Vesna Boraska Perica, Christopher S Franklin, James A B Floyd, Laura M Thornton, Laura M Huckins, Lorraine Southam, N William Rayner, Ioanna Tachmazidou, Kelly L Klump, Janet Treasure, Cathryn M Lewis, Ulrike Schmidt, Federica Tozzi, Kirsty Kiezebrink, Johannes Hebebrand, Philip Gorwood, Roger A H Adan, Martien J H Kas, Angela Favaro, Paolo Santonastaso, Fernando Fernández-Aranda, Monica Gratacos, Filip Rybakowski, Monika Dmitrzak-Weglarz, Jaakko Kaprio, Anna Keski-Rahkonen, Anu Raevuori-Helkamaa, Eric F Van Furth, Margarita C T Slof-Op’t Landt, James I Hudson, Ted Reichborn-Kjennerud, Gun Peggy S Knudsen, Palmiero Monteleone, Allan S Kaplan, Andreas Karwautz, Hakon Hakonarson, Wade H Berrettini, Yiran Guo, Dong Li, Nicholas J Schork, Gen Komaki, Tetsuya Ando, Hidetoshi Inoko, Tõnu Esko, Krista Fischer, Katrin Männik, Andres Metspalu, Jessica H Baker, Roger D Cone, Jennifer Dackor, Janiece E DeSocio, Christopher E Hilliard, Julie K O’Toole, Jacques Pantel, Jin P Szatkiewicz, Chrysecolla Taico, Stephanie Zerwas, Sara E Trace, Oliver S P Davis, Sietske Helder, Katharina Bühren, Roland Burghardt, Martina de Zwaan, Karin Egberts, Stefan Ehrlich, Beate Herpertz-Dahlmann, Wolfgang Herzog, Hartmut Imgart, André Scherag, Susann Scherag, Stephan Zipfel, Claudette Boni, Nicolas Ramoz, Audrey Versini, Marek K Brandys, Unna N Danner, Carolien de Kove, Judith Hendriks, Bobby P C Koeleman, Roel A Ophoff, Eric Strengman, Annemarie A van Elburg, Alice Bruson, Maurizio Clementi, Daniela Degortes, Monica Forzan, Elena Tenconi, Elisa Docampo, Geòrgia Escaramí, Susana Jiménez-Murcia, Jolanta Lissowska, Andrzej Rajewski, Neonila Szeszenia-Dabrowska, Agnieszka Slopien, Joanna Hauser, Leila Karhunen, Ingrid Meulenbelt, P Eline Slagboom, Alfonso Tortorella, Mario Maj, George Dedoussis, Dimitris Dikeos, Fragiskos Gonidakis, Konstantinos Tziouvas, Artemis Tsitsika, Hana Papezova, Lenka Slachtova, Debora Martaskova, James L Kennedy, Robert D Levitan, Zeynep Yilmaz, Julia Huemer, Doris Koubek, Elisabeth Merl, Gudrun Wagner, Paul Lichtenstein, Gerome Breen, Sarah Cohen-Woods, Anne Farmer, Peter McGuffin, Sven Cichon, Ina Giegling, Stefan Herms, Dan Rujescu, Stefan Schreiber, H-Erich Wichmann, Christian Dina, Rob Sladek, Giovanni Gambaro, Nicole Soranzo, Antonio Julia, Sara Marsal, Raquel Rabionet, Valerie Gaborieau, Danielle M Dick, Aarno Palotie, Samuli Ripatti, Elisabeth Widén, Ole A Andreassen, Thomas Espeseth, Astri Lundervold, Ivar Reinvang, Vidar M Steen, Stephanie Le Hellard, Morten Mattingsdal, Ioanna Ntalla, Vladimir Bencko, Lenka Foretova, Vladimir Janout, Marie Navratilova, Steven Gallinger, Dalila Pinto, Stephen W Scherer, Harald Aschauer, Laura Carlberg, Alexandra Schosser, Lars Alfredsson, Bo Ding, Lars Klareskog, Leonid Padyukov, Chris Finan, Gursharan Kalsi, Marion Roberts, Darren W Logan, Leena Peltonen, Graham R S Ritchie, Jeff C Barrett, Xavier Estivill, Anke Hinney, Patrick F Sullivan, David A Collier, Eleftheria Zeggini, and Cynthia M Bulik.

Wellcome Trust Case Control Consortium 3 (WTCCC3): Carl A Anderson, Jeffrey C Barrett, James A B Floyd, Christopher S Franklin, Ralph McGinnis, Nicole Soranzo, Eleftheria Zeggini, Jennifer Sambrook, Jonathan Stephens, Willem H Ouwehand, Wendy L McArdle, Susan M Ring, David P Strachan, Graeme Alexander, Cynthia M Bulik, David A Collier, Peter J Conlon, Anna Dominiczak, Audrey Duncanson, Adrian Hill, Cordelia Langford, Graham Lord, Alexander P Maxwell, Linda Morgan, Leena Peltonen, Richard N Sandford, Neil Sheerin, Frederik O Vannberg, Hannah Blackburn, Wei-Min Chen, Sarah Edkins, Mathew Gillman, Emma Gray, Sarah E Hunt, Suna Nengut-Gumuscu, Simon Potter, Stephen S Rich, Douglas Simpkin, and Pamela Whittaker.

The members of the ReproGen consortium are John RB Perry, Felix Day, Cathy E Elks, Patrick Sulem, Deborah J Thompson, Teresa Ferreira, Chunyan He, Daniel I Chasman, Tnu Esko, Gudmar Thorleifsson, Eva Albrecht, Wei Q Ang, Tanguy Corre, Diana L Cousminer, Bjarke Feenstra, Nora Franceschini, Andrea Ganna, Andrew D Johnson, Sanela Kjellqvist, Kathryn L Lunetta, George McMahon, Ilja M Nolte, Lavinia Paternoster, Eleonora Porcu, Albert V Smith, Lisette Stolk, Alexander Teumer, Natalia Ternikova, Emmi Tikkanen, Sheila Ulivi, Erin K Wagner, Najaf Amin, Laura J Bierut, Enda M Byrne, JoukeJan Hottenga, Daniel L Koller, Massimo Mangino, Tune H Pers, Laura M YergesArmstrong, Jing Hua Zhao, Irene L Andrulis, Hoda AntonCulver, Femke Atsma, Stefania Bandinelli, Matthias W Beckmann, Javier Benitez, Carl Blomqvist, Stig E Bojesen, Manjeet K Bolla, Bernardo Bonanni, Hiltrud Brauch, Hermann Brenner, Julie E Buring, Jenny ChangClaude, Stephen Chanock, Jinhui Chen, Georgia ChenevixTrench, J. Margriet Colle, Fergus J Couch, David Couper, Andrea D Coveillo, Angela Cox, Kamila Czene, Adamo Pio D’adamo, George Davey Smith, Immaculata De Vivo, Ellen W Demerath, Joe Dennis, Peter Devilee, Aida K Dieffenbach, Alison M Dunning, Gudny Eiriksdottir, Johan G Eriksson, Peter A Fasching, Luigi Ferrucci, Dieter FleschJanys, Henrik Flyger, Tatiana Foroud, Lude Franke, Melissa E Garcia, Montserrat GarcaClosas, Frank Geller, Eco EJ de Geus, Graham G Giles, Daniel F Gudbjartsson, Vilmundur Gudnason, Pascal Gunel, Suiqun Guo, Per Hall, Ute Hamann, Robin Haring, Catharina A Hartman, Andrew C Heath, Albert Hofman, Maartje J Hooning, John L Hopper, Frank B Hu, David J Hunter, David Karasik, Douglas P Kiel, Julia A Knight, VeliMatti Kosma, Zoltan Kutalik, Sandra Lai, Diether Lambrechts, Annika Lindblom, Reedik Mgi, Patrik K Magnusson, Arto Mannermaa, Nicholas G Martin, Gisli Masson, Patrick F McArdle, Wendy L McArdle, Mads Melbye Kyriaki Michailidou, Evelin Mihailov, Lili Milani, Roger L Milne, Heli Nevanlinna, Patrick Neven, Ellen A Nohr, Albertine J Oldehinkel, Ben A Oostra, Aarno Palotie,, Munro Peacock, Nancy L Pedersen, Paolo Peterlongo, Julian Peto, Paul DP Pharoah, Dirkje S Postma, Anneli Pouta, Katri Pylks, Paolo Radice, Susan Ring, Fernando Rivadeneira, Antonietta Robino, Lynda M Rose, Anja Rudolph, Veikko Salomaa, Serena Sanna, David Schlessinger, Marjanka K Schmidt, Mellissa C Southey, Ulla Sovio Meir J Stampfer, Doris Stckl Anna M Storniolo, Nicholas J Timpson Jonathan Tyrer, Jenny A Visser, Peter Vollenweider, Henry Vlzke, Gerard Waeber, Melanie Waldenberger, Henri Wallaschofski, Qin Wang, Gonneke Willemsen, Robert Winqvist, Bruce HR Wolffenbuttel, Margaret J Wright, Australian Ovarian Cancer Study The GENICA Network, kConFab, The LifeLines Cohort Study, The InterAct Consortium, Early Growth Genetics (EGG) Consortium, Dorret I Boomsma, Michael J Econs, KayTee Khaw, Ruth JF Loos, Mark I McCarthy, Grant W Montgomery, John P Rice, Elizabeth A Streeten, Unnur Thorsteinsdottir, Cornelia M van Duijn, Behrooz Z Alizadeh, Sven Bergmann, Eric Boerwinkle, Heather A Boyd, Laura Crisponi, Paolo Gasparini, Christian Gieger, Tamara B Harris, Erik Ingelsson, MarjoRiitta Jrvelin, Peter Kraft, Debbie Lawlor, Andres Metspalu, Craig E Pennell, Paul M Ridker, Harold Snieder, Thorkild IA Srensen, Tim D Spector, David P Strachan, Andr G Uitterlinden, Nicholas J Wareham, Elisabeth Widen, Marek Zygmunt, Anna Murray, Douglas F Easton, Kari Stefansson, Joanne M Murabito, Ken K Ong.

## Footnotes

1 We ignore the distinction between normalizing and centering in the population and in the sample, since this introduces only 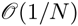 error.

2 The assumption that all *β* is drawn with equal variance for all SNPs hides an implicit assumption that rare SNPs have larger per-allele effect sizes than common SNPs. As discussed in the simulations section of the main text and in our earlier work [21], LD Score regression is robust to moderate violations of this assumption, though it may break down in extreme cases, *e.g.*, if all causal variants are rare. In situations where a different model for Var[*β*] is more appropriate, all proofs in this note go through with LD Score replaced by weighted LD Scores, 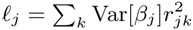.

3 For instance, it is sufficient but not necessary to assume that *β, γ, δ* and *ϵ* are multivariate normal. More generally, the *z*-scores will be approximately normal if *β* and *γ* are reasonably polygenic. If the distribution of effect sizes is heavy-tailed, *e.g.*, if there are few casual SNPs, then the CVF may be larger.

4 Conditional on the marginal effect of *j*, the expected value of 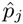 is not equal to *p_j_* unless *P* = *K* or the marginal effect of *j* is zero.

5 For *ℓ_j_* = 100 (roughly the median 1kG LD Score), *M* = 10^7^ and *ρ_g,obs_* = 1, we get *ρ_g,obs_ℓ_j_*/*M* = 10^−5^. A worst-case value for *N_s_*/*N*_1_*N*_2_ might be *N_s_* = *N*_1_ = *N*_2_ = 10^3^, in which case *N_s_*/*N*_1_*N*_2_ = 10^−3^. Thus, *ρ_g,obs_ℓ_j_*/*M* and *N_s_*/*N*_1_*N*_2_ will generally be at least 3 orders of magnitude smaller than 1.

## References

[1] George Davey Smith and Shah Ebrahim. Mendelian randomization: can genetic epidemiology contribute to understanding environmental determinants of disease? International journal of epidemiology, 32(1):1–22, 2003.

[2] George Davey Smith and Gibran Hemani. Mendelian randomization: genetic anchors for causal inference in epidemiological studies. Human molecular genetics, 23(R1):R89–R98, 2014.

[3] SG Vandenberg. Multivariate analysis of twin differences. Methods and goals in human behavior genetics, pages 29–43, 1965.

[4] Oscar Kempthorne and Richard H Osborne. The interpretation of twin data. American journal of human genetics, 13(3):320, 1961.

[5] John C Loehlin and Steven Gerritjan Vandenberg. Genetic and environmental components in the covariation of cognitive abilities: An additive model. Louisville Twin Study, University of Louisville, 1966.

[6] Michael Neale and Lon Cardon. Methodology for genetic studies of twins and families. Number 67. Springer, 1992.

[7] Paul Lichtenstein, Benjamin H Yip, Camilla Björk, Yudi Pawitan, Tyrone D Cannon, Patrick F Sullivan, and Christina M Hultman. Common genetic determinants of schizophrenia and bipolar disorder in swedish families: a population-based study. The Lancet, 373(9659):234–239, 2009.

[8] Joshua D Angrist and Jörn-Steffen Pischke. Mostly harmless econometrics: An empiricist’s companion. Princeton university press, 2008.

[9] Benjamin F Voight, Gina M Peloso, Marju Orho-Melander, Ruth Frikke-Schmidt, Maja Barbalic, Majken K Jensen, George Hindy, Hilma Hólm, Eric L Ding, Toby Johnson, et al. Plasma hdl cholesterol and risk of myocardial infarction: a mendelian randomisation study. The Lancet, 380(9841):572–580, 2012.

[10] Ron Do, Cristen J Willer, Ellen M Schmidt, Sebanti Sengupta, Chi Gao, Gina M Peloso, Stefan Gustafsson, Stavroula Kanoni, Andrea Ganna, Jin Chen, et al. Common variants associated with plasma triglycerides and risk for coronary artery disease. Nature genetics, 45(11):1345–1352, 2013.

[11] Peter M Visscher, Matthew A Brown, Mark I McCarthy, and Jian Yang. Five years of gwas discovery. The American Journal of Human Genetics, 90(1):7–24, 2012.

[12] Stephen Burgess, Simon G Thompson, et al. Avoiding bias from weak instruments in mendelian randomization studies. International journal of epidemiology, 40(3):755–764, 2011.

[13] Jian Yang, Beben Benyamin, Brian P McEvoy, Scott Gordon, Anjali K Henders, Dale R Nyholt, Pamela A Madden, Andrew C Heath, Nicholas G Martin, Grant W Montgomery, et al. Common snps explain a large proportion of the heritability for human height. Nature Genetics, 42(7):565–569, 2010.

[14] Jian Yang, S Hong Lee, Michael E Goddard, and Peter M Visscher. Gcta: a tool for genomewide complex trait analysis. The American Journal of Human Genetics, 88(1):76–82, 2011.

[15] Sang Hong Lee, Jian Yang, Michael E Goddard, Peter M Visscher, and Naomi R Wray. Estimation of pleiotropy between complex diseases using single-nucleotide polymorphism-derived genomic relationships and restricted maximum likelihood. Bioinformatics, 28(19):2540–2542, 2012.

[16] Cross-Disorder Group of the Psychiatric Genomics Consortium et al. Genetic relationship between five psychiatric disorders estimated from genome-wide snps. Nature Genetics, 2013.

[17] Shashaank Vattikuti, Juen Guo, and Carson C Chow. Heritability and genetic correlations explained by common snps for metabolic syndrome traits. PLoS genetics, 8(3):e1002637, 2012.

[18] Guo-Bo Chen, Sang Hong Lee, Marie-Jo A Brion, Grant W Montgomery, Naomi R Wray, Graham L Radford-Smith, Peter M Visscher, et al. Estimation and partitioning of (co) heritability of inflammatory bowel disease from gwas and immunochip data. Human molecular genetics, page ddu174, 2014.

[19] Shaun M Purcell, Naomi R Wray, Jennifer L Stone, Peter M Visscher, Michael C O’Donovan, Patrick F Sullivan, Pamela Sklar, Shaun M Purcell, Jennifer L Stone, Patrick F Sullivan, et al. Common polygenic variation contributes to risk of schizophrenia and bipolar disorder. Nature, 460(7256):748–752, 2009.

[20] Frank Dudbridge. Power and predictive accuracy of polygenic risk scores. PLoS genetics, 9(3):e1003348, 2013.

[21] Brendan Bulik-Sullivan, Po-Ru Loh, Hilary Finucane, Stephan Ripke, Jian Yang, Nick Patterson, Mark J Daly, Alkes L Price, and Benjamin M Neale. Ld score regression distinguishes confounding from polygenicity in genome-wide association studies. Nature Genetics, 2015.

[22] Jian Yang, Michael N Weedon, Shaun Purcell, Guillaume Lettre, Karol Estrada, Cristen J Willer, Albert V Smith, Erik Ingelsson, Jeffrey R O’Connell, Massimo Mangino, et al. Genomic inflation factors under polygenic inheritance. European Journal of Human Genetics, 19(7):807–812, 2011.

[23] Doug Speed, Gibran Hemani, Michael R Johnson, and David J Balding. Improved heritability estimation from genome-wide snps. The American Journal of Human Genetics, 91(6):1011–1021, 2012.

[24] Cross-Disorder Group of the Psychiatric Genomics Consortium et al. Identification of risk loci with shared effects on five major psychiatric disorders: a genome-wide analysis. Lancet, 381(9875):1371, 2013.

[25] John RB Perry, Felix Day, Cathy E Elks, Patrick Sulem, Deborah J Thompson, Teresa Ferreira, Chunyan He, Daniel I Chasman, Tõnu Esko, Gudmar Thorleifsson, et al. Parent-of-origin-specific allelic associations among 106 genomic loci for age at menarche. Nature, 514(7520):92–97, 2014.

[26] Andrew P Morris, Benjamin F Voight, Tanya M Teslovich, Teresa Ferreira, Ayellet V Segre, Valgerdur Steinthorsdottir, Rona J Strawbridge, Hassan Khan, Harald Grallert, Anubha Mahajan, et al. Large-scale association analysis provides insights into the genetic architecture and pathophysiology of type 2 diabetes. Nature genetics, 44(9):981, 2012.

[27] Momoko Horikoshi, Hanieh Yaghootkar, Dennis O Mook-Kanamori, Ulla Sovio, H Rob Taal, Branwen J Hennig, Jonathan P Bradfield, Beate St Pourcain, David M Evans, Pimphen Charoen, et al. New loci associated with birth weight identify genetic links between intrauterine growth and adult height and metabolism. Nature genetics, 45(1):76–82, 2013.

[28] Rachel M Freathy, Amanda J Bennett, Susan M Ring, Beverley Shields, Christopher J Groves, Nicholas J Timpson, Michael N Weedon, Eleftheria Zeggini, Cecilia M Lindgren, Hana Lango, et al. Type 2 diabetes risk alleles are associated with reduced size at birth. Diabetes, 58(6):1428–1433, 2009.

[29] Early Growth Genetics (EGG) Consortium et al. A genome-wide association meta-analysis identifies new childhood obesity loci. Nature genetics, 44(5):526–531, 2012.

[30] H Rob Taal, Beate St Pourcain, Elisabeth Thiering, Shikta Das, Dennis O Mook-Kanamori, Nicole M Warrington, Marika Kaakinen, Eskil Kreiner-Møller, Jonathan P Bradfield, Rachel M Freathy, et al. Common variants at 12q15 and 12q24 are associated with infant head circumference. Nature genetics, 44(5):532–538, 2012.

[31] NC Onland-Moret, PHM Peeters, CH Van Gils, F Clavel-Chapelon, T Key, A Tjønneland, A Trichopoulou, R Kaaks, Jonas Manjer, S Panico, et al. Age at menarche in relation to adult height the epic study. American journal of epidemiology, 162(7):623–632, 2005.

[32] Felix Day et al. Puberty timing associated with diabetes, cardiovascular disease and also diverse health outcomes in men and women: the uk biobank study. Submitted, 2014.

[33] Cathy E Elks, Ken K Ong, Robert A Scott, Yvonne T van der Schouw, Judith S Brand, Petra A Wark, Pilar Amiano, Beverley Balkau, Aurelio Barricarte, Heiner Boeing, et al. Age at menarche and type 2 diabetes risk the epic-interact study. Diabetes care, 36(11):3526–3534, 2013.

[34] Hilary K. Finucane, Brendan Bulik-Sullivan, Alexander Gusev, Gosia Trynka, Yakir Reshef, Po-Ru Loh, Verneri Anttila, Han Xu, Chongzhi Zang, Kyle Farh, Stephan Ripke, Felix R. Day, The ReproGen Consortium, Schizophrenia Working Group of the Psychiatric Genomics Consortium, Shaun Purcell, Eli Stahl, Sara Lindstrom, John R. B. Perry, Yukinori Okada, Brad Bernstein, Soumya Raychaudhuri, Mark Daly, Nick Patterson, Benjamin M. Neale, and Alkes L. Price. Polygenic effects of cell-type-specific functional elements in 17 traits and 1.3 million phenotyped samples. In preparation, 2014.

[35] I Sadaf Farooqi. Defining the neural basis of appetite and obesity: from genes to behaviour. Clinical Medicine, 14(3):286–289, 2014.

[36] Na Wang, Xianglan Zhang, Yong-Bing Xiang, Gong Yang, Hong-Lan Li, Jing Gao, Hui Cai, Yu-Tang Gao, Wei Zheng, and Xiao-Ou Shu. Associations of adult height and its components with mortality: a report from cohort studies of 135 000 chinese women and men. International journal of epidemiology, 40(6):1715–1726, 2011.

[37] Patricia R Hebert, Janet W Rich-Edwards, JE Manson, Paul M Ridker, Nancy R Cook, Gerald T O’Connor, Julie E Buring, and Charles H Hennekens. Height and incidence of cardiovascular disease in male physicians. Circulation, 88(4):1437–1443, 1993.

[38] Janet W Rich-Edwards, JoAnn E Manson, Meir J Stampfer, Graham A Colditz, Walter C Willett, Bernard Rosner, Frank E Speizer, and Charles H Hennekens. Height and the risk of cardiovascular disease in women. American journal of epidemiology, 142(9):909–917, 1995.

[39] Cornelius A Rietveld, Sarah E Medland, Jaime Derringer, Jian Yang, Tõnu Esko, Nicolas W Martin, Harm-Jan Westra, Konstantin Shakhbazov, Abdel Abdellaoui, Arpana Agrawal, et al. Gwas of 126,559 individuals identifies genetic variants associated with educational attainment. Science, 340(6139):1467–1471, 2013.

[40] Deborah E Barnes and Kristine Yaffe. The projected effect of risk factor reduction on alzheimer’s disease prevalence. The Lancet Neurology, 10(9):819–828, 2011.

[41] Sam Norton, Fiona E Matthews, Deborah E Barnes, Kristine Yaffe, and Carol Brayne. Potential for primary prevention of alzheimer’s disease: an analysis of population-based data. The Lancet Neurology, 13(8):788–794, 2014.

[42] James H MacCabe, Mats P Lambe, Sven Cnattingius, Pak C Sham, Anthony S David, Abraham Reichenberg, Robin M Murray, and Christina M Hultman. Excellent school performance at age 16 and risk of adult bipolar disorder: national cohort study. The British Journal of Psychiatry, 196(2):109–115, 2010.

[43] Jari Tiihonen, Jari Haukka, Markus Henriksson, Mary Cannon, Tuula Kieseppä, Ilmo Laaksonen, Juhani Sinivuo, and Jouko Lönnqvist. Premorbid intellectual functioning in bipolar disorder and schizophrenia: results from a cohort study of male conscripts. American Journal of Psychiatry, 162(10):1904–1910, 2005.

[44] John P Pierce, Michael C Fiore, Thomas E Novotny, Evridiki J Hatziandreu, and Ronald M Davis. Trends in cigarette smoking in the united states: educational differences are increasing. Jama, 261(1):56–60, 1989.

[45] Ruth H Striegel-Moore, Vicki Garvin, Faith-Anne Dohm, and Robert A Rosenheck. Psychiatric comorbidity of eating disorders in men: a national study of hospitalized veterans. International Journal of Eating Disorders, 25(4):399–404, 1999.

[46] Barton J Blinder, Edward J Cumella, and Visant A Sanathara. Psychiatric comorbidities of female inpatients with eating disorders. Psychosomatic Medicine, 68(3):454–462, 2006.

[47] Ian J Deary, Steve Strand, Pauline Smith, and Cres Fernandes. Intelligence and educational achievement. Intelligence, 35(1):13–21, 2007.

[48] Catherine M Calvin, Cres Fernandes, Pauline Smith, Peter M Visscher, and Ian J Deary. Sex, intelligence and educational achievement in a national cohort of over 175,000 11-year-old schoolchildren in england. Intelligence, 38(4):424–432, 2010.

[49] Maureen S Durkin, Matthew J Maenner, F John Meaney, Susan E Levy, Carolyn DiGuiseppi, Joyce S Nicholas, Russell S Kirby, Jennifer A Pinto-Martin, and Laura A Schieve. Socioeconomic inequality in the prevalence of autism spectrum disorder: evidence from a us cross-sectional study. PLoS One, 5(7):e11551, 2010.

[50] Elise B Robinson, Kaitlin E Samocha, Jack A Kosmicki, Lauren McGrath, Benjamin M Neale, Roy H Perlis, and Mark J Daly. Autism spectrum disorder severity reflects the average contribution of de novo and familial influences. Proceedings of the National Academy of Sciences, 111(42):15161–15165, 2014.

[51] Kaitlin E Samocha, Elise B Robinson, Stephan J Sanders, Christine Stevens, Aniko Sabo, Lauren M McGrath, Jack A Kosmicki, Karola Rehnström, Swapan Mallick, Andrew Kirby, et al. A framework for the interpretation of de novo mutation in human disease. Nature genetics, 46(9):944–950, 2014.

[52] Alan J Silman and Jacqueline E Pearson. Epidemiology and genetics of rheumatoid arthritis. Arthritis Res, 4(Suppl 3):S265–S272, 2002.

[53] Jose de Leon and Francisco J Diaz. A meta-analysis of worldwide studies demonstrates an association between schizophrenia and tobacco smoking behaviors. Schizophrenia research, 76(2):135–157, 2005.

[54] Ole A Andreassen, Srdjan Djurovic, Wesley K Thompson, Andrew J Schork, Kenneth S Kendler, Michael C O?Donovan, Dan Rujescu, Thomas Werge, Martijn van de Bunt, Andrew P Morris, et al. Improved detection of common variants associated with schizophrenia by leveraging pleiotropy with cardiovascular-disease risk factors. The American Journal of Human Genetics, 92(2):197–209, 2013.

[55] Chris Cotsapas, Benjamin F Voight, Elizabeth Rossin, Kasper Lage, Benjamin M Neale, Chris Wallace, Gonçalo R Abecasis, Jeffrey C Barrett, Timothy Behrens, Judy Cho, et al. Pervasive sharing of genetic effects in autoimmune disease. PLoS genetics, 7(8):e1002254, 2011.

[56] Kyle Kai-How Farh, Alexander Marson, Jiang Zhu, Markus Kleinewietfeld, William J Housley, Samantha Beik, Noam Shoresh, Holly Whitton, Russell JH Ryan, Alexander A Shishkin, et al. Genetic and epigenetic fine mapping of causal autoimmune disease variants. Nature, 2014.

[57] Peter Wurtz et al. Metabolic signatures of adiposity in young adults: Mendelian randomization analysis and effects of weight change. PLoS Medicine, 2014.

[58] Stephen Burgess, Daniel F Freitag, Hassan Khan, Donal N Gorman, and Simon G Thompson. Using multivariable mendelian randomization to disentangle the causal effects of lipid fractions. PloS one, 9(10):e108891, 2014.

[59] Sander Greenland, Judea Pearl, and James M Robins. Causal diagrams for epidemiologic research. Epidemiology, pages 37–48, 1999.

[60] Andy Dahl, Victoria Hore, Valentina Iotchkova, and Jonathan Marchini. Network inference in matrix-variate gaussian models with non-independent noise. arXiv preprint arXiv:1312.1622, 2013.

[61] Hugues Aschard, Bjarni J Vilhjálmsson, Amit D Joshi, Alkes L Price, and Peter Kraft. Adjusting for heritable covariates can bias effect estimates in genome-wide association studies. The American Journal of Human Genetics, 2015.

[62] International HapMap 3 Consortium et al. Integrating common and rare genetic variation in diverse human populations. Nature, 467(7311):52–58, 2010.

[63] Karl Pearson and Alice Lee. On the inheritance of characters not capable of exact quantitative measurement. Philosophical Transactions of the Royal Society of London, A (195), pages 79–150, 1901.

[64] Schizophrenia Working Group of the Psychiatric Genomics Consortium et al. Biological insights from 108 schizophrenia-associated genetic loci. Nature, 511(7510):421–427, 2014.

[65] Pamela Sklar, Stephan Ripke, Laura J Scott, Ole A Andreassen, Sven Cichon, Nick Craddock, Howard J Edenberg, John I Nurnberger, Marcella Rietschel, Douglas Blackwood, et al. Large-scale genome-wide association analysis of bipolar disorder identifies a new susceptibility locus near odz4. Nature genetics, 43(10):977, 2011.

[66] Stephan Ripke, Naomi R Wray, Cathryn M Lewis, Steven P Hamilton, Myrna M Weissman, Gerome Breen, Enda M Byrne, Douglas HR Blackwood, Dorret I Boomsma, Sven Cichon, et al. A mega-analysis of genome-wide association studies for major depressive disorder. Molecular psychiatry, 18(4):497–511, 2012.

[67] Vesna Boraska, Christopher S Franklin, James AB Floyd, Laura M Thornton, Laura M Huck-ins, Lorraine Southam, N William Rayner, Ioanna Tachmazidou, Kelly L Klump, Janet Treasure, et al. A genome-wide association study of anorexia nervosa. Molecular psychiatry, 2014.

[68] Tobacco, Genetics Consortium et al. Genome-wide meta-analyses identify multiple loci associated with smoking behavior. Nature genetics, 42(5):441–447, 2010.

[69] Jean-Charles Lambert, Carla A Ibrahim-Verbaas, Denise Harold, Adam C Naj, Rebecca Sims, Céline Bellenguez, Gyungah Jun, Anita L DeStefano, Joshua C Bis, Gary W Beecham, et al. Meta-analysis of 74,046 individuals identifies 11 new susceptibility loci for alzheimer’s disease. Nature genetics, 2013.

[70] Hana Lango Allen, Karol Estrada, Guillaume Lettre, Sonja I Berndt, Michael N Weedon, Fernando Rivadeneira, Cristen J Willer, Anne U Jackson, Sailaja Vedantam, Soumya Raychaudhuri, et al. Hundreds of variants clustered in genomic loci and biological pathways affect human height. Nature, 467(7317):832–838, 2010.

[71] Sonja I Berndt, Stefan Gustafsson, Reedik Mägi, Andrea Ganna, Eleanor Wheeler, Mary F Feitosa, Anne E Justice, Keri L Monda, Damien C Croteau-Chonka, Felix R Day, et al. Genome-wide meta-analysis identifies 11 new loci for anthropometric traits and provides insights into genetic architecture. Nature genetics, 45(5):501–512, 2013.

[72] Heribert Schunkert, Inke R König, Sekar Kathiresan, Muredach P Reilly, Themistocles L Assimes, Hilma Holm, Michael Preuss, Alexandre FR Stewart, Maja Barbalic, Christian Gieger, et al. Large-scale association analysis identifies 13 new susceptibility loci for coronary artery disease. Nature genetics, 43(4):333–338, 2011.

[73] Tanya M Teslovich, Kiran Musunuru, Albert V Smith, Andrew C Edmondson, Ioannis M Stylianou, Masahiro Koseki, James P Pirruccello, Samuli Ripatti, Daniel I Chasman, Cristen J Willer, et al. Biological, clinical and population relevance of 95 loci for blood lipids. Nature, 466(7307):707–713, 2010.

[74] Alisa K Manning, Marie-France Hivert, Robert A Scott, Jonna L Grimsby, Nabila Bouatia-Naji, Han Chen, Denis Rybin, Ching-Ti Liu, Lawrence F Bielak, Inga Prokopenko, et al. A genome-wide approach accounting for body mass index identifies genetic variants influencing fasting glycemic traits and insulin resistance. Nature genetics, 44(6):659–669, 2012.

[75] Ralf JP van der Valk, Eskil Kreiner-Møller, Marjolein N Kooijman, Mònica Guxens, Evangelia Stergiakouli, Annika Sääf, Jonathan P Bradfield, Frank Geller, M Geoffrey Hayes, Diana L Cousminer, et al. A novel common variant in dcst2 is associated with length in early life and height in adulthood. Human molecular genetics, page ddu510, 2014.

[76] Luke Jostins, Stephan Ripke, Rinse K Weersma, Richard H Duerr, Dermot P McGovern, Ken Y Hui, James C Lee, L Philip Schumm, Yashoda Sharma, Carl A Anderson, et al. Host-microbe interactions have shaped the genetic architecture of inflammatory bowel disease. Nature, 491(7422):119–124, 2012.

[77] Eli A Stahl, Soumya Raychaudhuri, Elaine F Remmers, Gang Xie, Stephen Eyre, Brian P Thomson, Yonghong Li, Fina AS Kurreeman, Alexandra Zhernakova, Anne Hinks, et al. Genome-wide association study meta-analysis identifies seven new rheumatoid arthritis risk loci. Nature genetics, 42(6):508–514, 2010.

[78] Schizophrenia Psychiatric Genome-Wide Association Study (GWAS) Consortium et al. Genome-wide association study identifies five new schizophrenia loci. Nature genetics, 43(10):969–976, 2011.

